# Two isoforms of the essential *C. elegans* Argonaute CSR-1 differentially regulate sperm and oocyte fertility through distinct small RNA classes

**DOI:** 10.1101/2020.07.20.212050

**Authors:** Amanda G. Charlesworth, Nicolas J. Lehrbach, Uri Seroussi, Mathias S. Renaud, Ruxandra I. Molnar, Jenna R. Woock, Matthew J. Aber, Annette J. Diao, Gary Ruvkun, Julie M. Claycomb

## Abstract

The *C. elegans* genome encodes nineteen functional Argonaute proteins that utilize 22G-RNAs, 26G-RNAs, miRNAs, or piRNAs to regulate their target transcripts. Only one of these proteins is essential under normal laboratory conditions: CSR-1. While CSR-1 has been studied in various developmental and functional contexts, nearly all studies investigating CSR-1 have overlooked the fact that the *csr-1* locus encodes two isoforms. These isoforms differ by an additional 163 amino acids present in the N-terminus of CSR-1a. Using CRISPR-Cas9 genome editing to introduce GFP::3xFLAG epitopes into the long (CSR-1a) and short (CSR-1b) isoforms of CSR-1, we identified differential expression patterns for the two isoforms. CSR-1a is expressed specifically during spermatogenesis and in select somatic tissues, including the intestine. In contrast, CSR-1b, is expressed constitutively in the germline. Essential functions of *csr-1* described in the literature coincide with CSR-1b. In contrast, CSR-1a plays tissue specific functions during spermatogenesis, where it integrates into a spermatogenesis sRNA regulatory network including ALG-3, ALG-4, and WAGO-10 that is necessary for male fertility. CSR-1a is also required in the intestine for the silencing of repetitive transgenes. Sequencing of small RNAs associated with each CSR-1 isoform reveals that CSR-1a engages with 22G- and 26G-RNAs, while CSR-1b interacts with only 22G-RNAs to regulate distinct groups of germline genes and regulate both sperm and oocyte-mediated fertility.

## INTRODUCTION

From the discovery of miRNAs to understanding the molecular mechanisms of RNA interference (RNAi), the nematode *C. elegans* has been a champion of sRNA biology (Youngman and Claycomb, 2014). At the heart of small RNA (sRNA) mediated gene regulatory pathways are the conserved Argonaute (AGO) proteins, which are guided in a sequence specific manner by sRNAs to impart a variety of gene regulatory outcomes on target transcripts, ranging from transcriptional modulation to RNA turnover and translational inhibition (Meister, 2013). Like most nematodes, *C. elegans* possesses a remarkable expanded family of nineteen AGO proteins which function in association with four types of sRNAs (including miRNAs, piRNAs, 22G-RNAs, and 26G-RNAs) to orchestrate normal developmental and differentiation programs as well as to navigate stressful environmental conditions, such as heat stress (Yigit et al., 2006) (Seroussi in preparation). Moreover, the scope of sRNA mediated gene regulatory potential in the worm is even more complex when considering that several AGOs, including ALG-1, ALG-2, ERGO-1, PPW-1, and CSR-1 are predicted to be expressed as multiple isoforms. Furthermore, many *C. elegans* AGOs have not been characterized, and even for those which have been studied, the distinct and overlapping functions of different isoforms remain almost entirely unexplored (Youngman and Claycomb, 2014).

The AGO CSR-1 (Chromosome Segregation and RNAi Deficient) is one of the most intensively studied and fascinating AGOs in *C. elegans*, owing in part to the fact that it is the only solely essential *ago*, and in part to its potentially conflicting and varied roles in gene regulation throughout development (Yigit et al., 2006; Claycomb et al., 2009). The *csr-1* locus encodes two isoforms, CSR-1a and CSR-1b, that vary only in their N-terminus by 163 amino acids (Claycomb et al., 2009). Within this extended N-terminal region of CSR-1a are fifteen individual arginine/glycine RGG/RG motifs, many of which are arranged into di- or tri-RGG/RG motifs (Thandapani et al., 2013). In other AGOs, including Drosophila Piwi, RGG/RG motifs are the sites of arginine di-methylation, which is required for the recruitment of additional factors, such as Tudor proteins, to ensure functionality of the AGO/sRNA complex (called the RISC; RNA Induced Silencing Complex) (Kirino et al., 2009; Liu et al., 2010; Vagin et al., 2009). These motifs are also typical of RNA binding proteins, and have been identified in association with phase separated RNA rich cytoplasmic granules, such as germ granules (Chong et al., 2018). The role and importance of CSR-1’s RGG/RG motifs have yet to be explored. Furthermore, CSR-1 has been widely studied as a single protein, thus any differential functions of CSR-1 isoforms during development are currently unknown.

CSR-1 is expressed in both the germline and soma in hermaphrodites (Claycomb et al., 2009; Avgousti et al., 2012). Immunostaining with antibodies that recognize both CSR-1 isoforms showed that CSR-1 is expressed in the hermaphrodite germline throughout all stages of germline development, from the two-cell embryo to spermatogenesis and oogenesis (Claycomb et al., 2009). Within the germline, CSR-1 is present in the cytoplasm, and is enriched in phase separated germ granules known as P granules (Claycomb et al., 2009). In oocytes and during early embryonic cell divisions, CSR-1 is present within the nucleus and associates with condensed chromosomes upon nuclear envelope breakdown (Claycomb et al., 2009). In the soma, CSR-1 was observed within intestinal cells, where it was detected in both the cytoplasm and nucleus (Claycomb et al., 2009; Avgousti et al., 2012). The germline expression pattern of CSR-1 highlights its role in fertility, as homozygous null mutants that eliminate the expression of both *csr-1* isoforms are nearly sterile, and the few embryos produced display chromosome segregation defects during early embryonic divisions (Yigit et al., 2006; Claycomb et al., 2009; Gerson-Gurwitz et al., 2016). However, which CSR-1 isoform is expressed in which tissues, and whether either isoform or both are necessary for fertility remain open questions.

While most AGOs are thought of as gene silencers, CSR-1 has been shown to play both positive (licensing) and negative (silencing) gene regulatory roles (Wedeles et al., 2013a). CSR-1 associates with a type of endogenous sRNA known as 22G-RNAs that target nearly 5000 germline-expressed protein coding genes (about 25% of the genome) (Conine et al., 2013; Claycomb et al., 2009). CSR-1-associated 22G-RNAs are produced by the RNA dependent RNA polymerase (RdRP), EGO-1, one of four RdRPs present in the worm (Claycomb et al., 2009; Gu et al., 2009). Microarray and GRO-seq studies demonstrated that loss of *csr-1* does not result in the global up-regulation of the majority of target transcripts (Conine et al., 2013; Claycomb et al., 2009; Cecere et al., 2014; Gerson-Gurwitz et al., 2016). Instead most CSR-1 target transcripts decrease in expression levels in *csr-1* mutants, pointing to its role in the protection or licensing of target transcripts, rather than their destruction or silencing.

There are several ways in which CSR-1 has been implicated in promoting the expression of its target transcripts. First, CSR-1 has been shown to promote the expression of histones. In this capacity, CSR-1 is thought to act as an endonuclease or “Slicer” (see below), where it promotes the processing of histone mRNAs by cleaving the 3′ stem loop of histone RNA transcripts enabling them to be translated. Loss of *csr-1* leads to decreased levels of histone transcripts and proteins (Avgousti et al., 2012). Second, CSR-1 licenses the expression of germline GFP transgenes in opposition to the silencing effects of the piRNA pathway, in a process related to paramutation, termed RNA activation (RNAa) (Seth et al., 2013; Wedeles et al., 2013b). CSR-1 also promotes the formation of euchromatin at its numerous target gene loci throughout the genome, and loss of *csr-1* leads to aberrant accumulation of heterochromatic histone modifications and decreased transcription at target gene loci (Gushchanskaia et al., 2019; Cecere et al., 2014).

CSR-1 is one of only a few AGOs (ALG-1, ALG-2, RDE-1, ERGO-1) which possess a predicted Slicer motif, DEDH, within its PIWI domain, rendering it capable of endonucleolytic cleavage of target transcripts. CSR-1 exhibits Slicer activity *in vitro* (Aoki et al., 2007), and mutation of these residues leads to embryonic lethality, although at a later developmental stage than null mutants (Gerson-Gurwitz et al., 2016). This Slicer activity is required for fine tuning gene expression during embryogenesis, as subsequent studies have linked embryonic chromosome segregation defects not to a direct role for CSR-1 in organizing chromatin, but to up-regulation of critical kinetochore and cell cycle components which are targeted by the CSR-1 sRNA pathway in the early embryo (Gerson-Gurwitz et al., 2016; Claycomb et al., 2009). Moreover, CSR-1’s Slicer activity is also thought to be important for RNAi, processing histone mRNA transcripts, and clearing maternal mRNAs at the oocyte to embryo transition (Aoki et al., 2007; Avgousti et al., 2012; Fassnacht et al., 2018).

While most studies of CSR-1 have focused on the oogenic gonad and embryos, CSR-1 has also been implicated in spermatogenesis. Spermatogenesis occurs in the L4 larval stage of development, prior to the onset of oogenesis (L’Hernault, 2006). In spermatogenesis, ALG-3 and −4 regulate a set of spermatogenesis transcripts both positively and negatively via a different set of endogenous sRNA called 26G-RNAs (Conine et al., 2013; Conine et al., 2010). 26G-RNAs are produced by the RdRP RRF-3, which makes long dsRNA that is processed by the endonuclease Dicer. The targeting of transcripts by 26G-RNAs leads to the amplification of 22G-RNAs by the RdRPs EGO-1 and RRF-1, and the subsequent targeting of the same transcripts by different 22G-RNA/AGO RISCs (Han et al., 2009; Conine et al., 2010; Gent et al., 2009; Vasale et al., 2010). Loss of *alg-3* and *alg-4* together leads to a progressive loss of fertility (Conine et al., 2013; Conine et al., 2010). Curiously, while loss of *csr-1* leads to sterile hermaphrodites (Yigit et al., 2006; Claycomb et al., 2009), *csr-1* null mutant males show a progressive loss of fertility similar to that of *alg-3, alg-4* double mutants (Conine et al., 2013).

The characterization of CSR-1 associated 22G-RNAs in males showed that CSR-1 targets a set of protein coding genes in common with CSR-1 targets in hermaphrodites undergoing oogenesis, along with an additional set of CSR-1 target genes. Furthermore, a subset of the genes targeted by CSR-1 during spermatogenesis overlap with genes targeted by ALG-3 and −4. Because 26G-RNAs associated with ALG-3 and −4 are thought to lead to the production of 22G-RNAs, this places CSR-1 downstream of ALG-3 and −4 during spermatogenesis (Conine et al., 2013). Consistent with its role in oogenesis, CSR-1 positively regulates its spermatogenic targets at the level of transcription during spermatogenesis (Conine et al., 2013), and depletion of *csr-1* by RNAi leads to the upregulation of spermatogenesis genes in oogenesis (Campbell and Updike, 2015). Why CSR-1 regulates the same subsets of genes in the spermatogenic and oogenic gonad, and which CSR-1 isoform(s) affect male fertility remain to be determined. In addition, transcriptomic analyses of isolated male and female *C. elegans* gonads showed that the *csr-1a* mRNA is abundant in the spermatogenic gonad, but is not highly expressed during oogenesis, pointing to a potential role for CSR-1a specifically in the spermatogenic germline that has not been explored (Ortiz et al., 2014).

Here we show that the two isoforms of CSR-1 have distinct expression profiles throughout development. The long isoform, CSR-1a, is expressed only during spermatogenesis within the germline, while the short isoform, CSR-1b, is expressed constitutively in the germline throughout all stages of development. During spermatogenesis, CSR-1a and CSR-1b associate with distinct subsets of sRNAs to target different sets of germline genes. While CSR-1b associates with 22G-RNAs to regulate gender neutral (i.e. expressed both during spermatogenesis and oogenesis) and oogenic transcripts (Ortiz et al., 2014), CSR-1a associates with both 26G-RNAs and 22G-RNAs to target spermatogenesis enriched transcripts (Ortiz et al., 2014). CSR-1a and another uncharacterized AGO, WAGO-10 integrate into the ALG-3 and −4 26G-RNA pathway to enable male fertility under elevated temperature, while CSR-1b is essential for hermaphrodite fertility. CSR-1a is also expressed in several somatic tissues, including the intestine, where it plays a role in silencing somatic transgenes. Collectively, our data indicate that different AGO isoforms are capable of vastly different functions and highlight the importance of studying each isoform as a separate entity.

## RESULTS

### CSR-1 isoforms have distinct expression patterns throughout development

The c*sr-1* locus encodes two isoforms that vary by the addition of 163 amino acids to the N-terminal region of CSR-1a. Several previous reports suggested that the two isoforms of CSR-1 (Fig. 1A) were differentially expressed throughout development (Ortiz et al., 2014; Claycomb et al., 2009), therefore we set out to examine and characterize any independent roles of the CSR-1 isoforms. To do so, we generated an extensive set of *csr-1* mutants and endogenously tagged strains using CRISPR-Cas9 genome editing (Dickinson et al., 2015; Kim et al., 2014) and MiniMos single copy transgene insertions (Frøkjær-Jensen et al., 2014) (Fig. S1A). We introduced GFP::3xFLAG into the N-terminus of the long isoform, CSR-1a, via CRISPR-Cas9 genome editing at the endogenous *csr-1* locus (Dickinson et al., 2015). Because the *csr-1b* locus is entirely shared with the *csr-1a* locus, epitope tagging CSR-1b invariably results in the internal epitope tagging of CSR-1a, thus we used this to our advantage to epitope tag both isoforms in a single strain (Ouyang et al., 2019). However, having both isoforms tagged precludes our ability to study CSR-1b separately. Therefore, we also introduced a stop codon into the *csr-1a* specific first exon, to generate a third strain possessing a null mutant for *csr-1a*, and expressing only GFP::3xFLAG::CSR-1b (referred to as *csr-1a(tor159)* in figures where *csr-1a* loss of function is assessed). To determine if the epitope tagged versions of CSR-1 were functional, we assessed the viable brood size of each strain, and found that all strains were fully fertile under normal lab culture conditions of 20°C (Fig. S1B).

**Figure 1.**
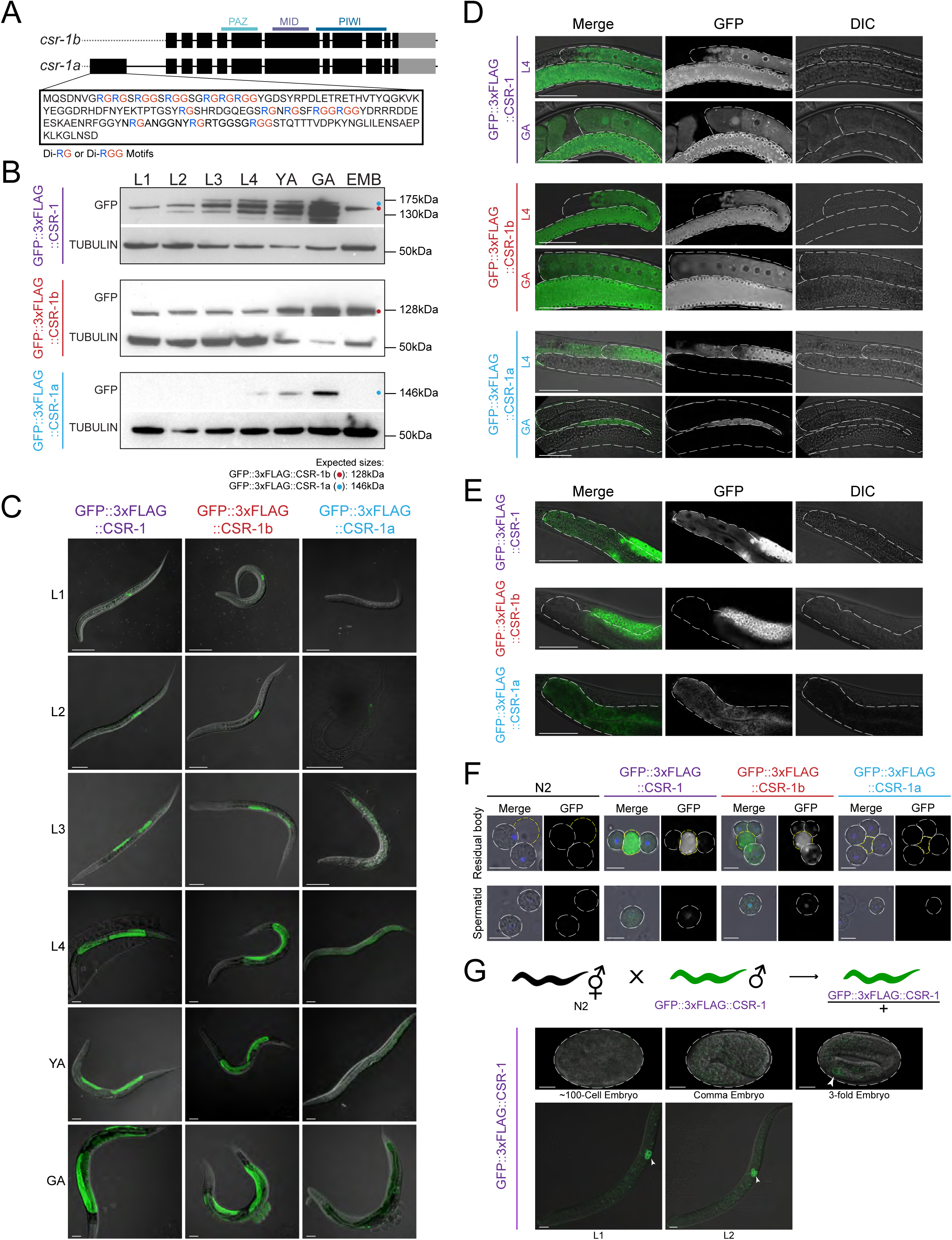
CSR-1a and CSR-1b isoforms are differentially expressed throughout development. **A)** Schematic representation of CSR-1 isoforms. Black boxes are exons, gray boxes are 3′UTRs. **B)** Western blot using GFP antibodies to detect GFP::3xFLAG::CSR-1, GFP::3xFLAG::CSR-1b, and GFP::3xFLAG::CSR-1a expression in each life stage (larval stages 1 to 4 (L1-L4), young adult (YA), gravid adult (GA) and embryo (EMB) stages). **C)** Fluorescence micrographs of whole animals expressing endogenously tagged CSR-1 isoforms during each life stage (L1-L4, YA and GA). Scale bar, 50μm. **D)** Fluorescence micrographs of animals in L4 and GA stages expressing GFP::3xFLAG::CSR-1, GFP::3xFLAG::CSR-1b and GFP::3xFLAG::CSR-1a in the germlines. Scale bar, 50μm. **E)** Fluorescence micrographs of animals expressing GFP::3xFLAG::CSR-1, GFP::3xFLAG::CSR-1b and GFP::3xFLAG::CSR-1a in the intestine. Scale bar, 50μm. **F)** Fluorescence micrographs of residual bodies (outlined in yellow) and spermatids (outlined in white) from animals expressing GFP::3xFLAG::CSR-1, GFP::3xFLAG::CSR-1b and GFP::3xFLAG::CSR-1a and stained with DAPI (blue in Merge), compared to wild-type (N2) animals. Scale bar, 5μm. **G)** Fluorescence micrographs showing zygotic expression of GFP::3xFLAG::CSR-1 in embryos, and L1 and L2 animals. Embryos were generated from crosses of wild-type hermaphrodites to *gfp::3xflag::csr-1* males. Scale bar, 10μm.

Next, we evaluated the tissues and developmental time points at which each isoform is expressed using western blotting (Fig. 1B) and confocal microscopy (Fig. 1C-E, Fig. S1C, Fig. S2, Fig. S3). In these experiments, CSR-1a and CSR-1b displayed clear differences in the timing and tissue-specificity of expression. Western blotting on the GFP::3xFLAG::CSR-1b strain demonstrated that, consistent with published data, CSR-1b is expressed during all developmental stages in hermaphrodites, with the highest levels of protein present in adults and in embryos (Fig. 1B). In examining this strain by confocal microscopy, we observed that CSR-1b is constitutively present in the germ cells and germline at all developmental stages in hermaphrodites, including the primordial germ cells in embryos, the spermatogenic gonad, and the oogenic gonad (Fig. 1C, Fig. S2). Consistent with expression during spermatogenesis in hermaphrodites (Fig. 1D), CSR-1b is also expressed in the male gonad (Fig. S2). Within all germline tissues, CSR-1b is present primarily in the cytoplasm and robustly localizes to germ granules, including what are likely to be P granules (Fig. 1D, S2, data not shown), and the newly identified Z granules (Fig. S1E) (Wan et al., 2018).

Because CSR-1b is expressed in both the spermatogenic and oogenic gonad (Fig. 1D), we next asked whether the protein was transmitted to progeny via sperm or oocytes. First, we performed confocal microscopy on sperm released from males. We observed that much of the CSR-1b pool was present in the residual body, a structure where sperm cytoplasmic contents are discarded during spermatogenesis (Fig. 1F) (L’Hernault, 2006). We also detected CSR-1b in mature spermatids, where it was present both in the nucleus and cytoplasmic foci (Fig. 1F) (Conine et al., 2013). These observations indicate that CSR-1b has the potential to be passed from males to progeny via sperm. In oocytes, CSR-1b localizes to the cytoplasm, germ granules, and oocyte chromatin (Fig. 1D, Fig. S1D,E), and could be maternally provisioned (Claycomb et al., 2009; Chu et al., 2006). In self-fertilizing GFP::3xFLAG::CSR-1b hermaphrodites, we observed GFP expression in early embryos, prior to the onset of zygotic transcription (Fig. S1C). In contrast, we did not observe GFP expression in early embryos produced from crosses between GFP::3xFLAG::CSR-1 males to wild-type hermaphrodites (Fig. 1G, and see below). These data indicate that CSR-1b is maternally inherited.

During early embryogenesis until approximately the 50 cell stage, maternally provisioned CSR-1b is detected in both somatic and germ cells (Fig. S1C, S2). In both cell types, CSR-1b is present in the cytoplasm and localizes to the nucleus in a cell-cycle dependent manner, coincident with mitosis (Fig. S1C, S2). In the germ cell lineage, CSR-1b also localizes to germ granules constitutively at all stages of germ cell development (Fig. 1C,D). To determine when zygotic expression of CSR-1b initiates, we crossed GFP::3xFLAG::CSR-1 males to wild-type hermaphrodites, and first detected GFP::3xFLAG::CSR-1 expression in the primordial germ cells Z2 and Z3 (Fig. 1G). [Note, although we used CSR-1 (total) in this experiment, we infer that this zygotic expression in Z2/Z3 is attributable to CSR-1b because CSR-1a is not maternally deposited nor expressed zygotically until the L2 stage (Fig. 1C, and see below).] Collectively, these data indicate that CSR-1b is maternally, and potentially paternally inherited, and that zygotic expression of CSR-1b initiates when the primordial germ cells become transcriptionally active during late embryonic development.

In contrast, western blotting demonstrated that CSR-1a displays a more restricted expression profile, and is detected only in L4, young adult, and gravid adult developmental stages (Fig. 1B). Confocal imaging throughout all developmental stages revealed that GFP::3xFLAG::CSR-1a is expressed in the germline only during the L4 larval stage in hermaphrodites, during which sperm are produced (Fig. 1C-D, S3). At this time it is also present in several somatic tissues, including the intestine (see below, Fig. 1E, S3). In the spermatogenic gonad, CSR-1a is present in the cytoplasm and within germ granules (Fig. 1D). We next examined dissected sperm to determine if CSR-1a could be passed to progeny paternally and found that CSR-1a is not packaged into mature spermatids, nor is it segregated into the residual body (Fig. 1F). As anticipated, CSR-1a is also expressed in the male gonad similarly to its expression in L4 hermaphrodites (Fig. S3). Because we do not observe CSR-1a in sperm or oocytes, we conclude that CSR-1a is not transmitted paternally or maternally to progeny.

In adult hermaphrodites, we were unable to detect CSR-1a in the oogenic germline, including within oocytes (Fig. 1D, S1D). Consistent with this, we did not observe the presence of CSR-1a in embryos at any stage (Fig. S1C). Nevertheless, western blotting data demonstrated the presence of CSR-1a in adults (Fig. 1B), which suggested that CSR-1a may be expressed in the soma. Indeed, we observed CSR-1a in several adult somatic tissues of both hermaphrodites and males (Fig. S3), including the intestine (Fig. 1E), spermatheca (Fig. S3), and several cells within the tail (Fig. S3). In adult males, CSR-1a is also present in the vas deferens and seminal vesicle (Fig. S3). When we looked at CSR-1a expression in early developmental stages, we observed its expression in the intestine from the L3 stage onward (Fig. 1C). In contrast, CSR-1b was not observed in any of these somatic tissues. Consistent with this intestinal expression pattern, using existing transcription factor binding data (Kudron et al., 2018), we observed enrichment of the intestinal master transcription factor ELT-2 (Fukushige et al., 1998) upstream of the *csr-1a* transcription start site at the L3 stage of development (Fig. S1F). Within the first intron of the *csr-1* locus are potential regulatory elements for *csr-1b*. Within this intron, we did not observe ELT-2 binding, but instead observed enrichment of several germline enriched transcription factors, including members of the SNAP-C (Kasper et al., 2014) and DRM complexes (Harrison et al., 2006), which we predict would regulate the expression of *csr-1b* in the germline. Taken together, these results highlight the potential for CSR-1a to play novel roles during spermatogenesis and within specific somatic tissues, including the intestine.

### CSR-1b is the essential CSR-1 isoform but CSR-1a can compensate

*csr-1* is the only singly essential *ago* in *C. elegans*. Homozygous zygotic null mutants of *csr-1* (which affect both isoforms) that are generated from heterozygous hermaphrodites develop to adulthood, but display defects in oogenesis. Consequently, they produce few embryos, and those few embryos that are generated display chromosome segregation defects and arrest early in embryogenesis, prior to the 50 cell stage (Claycomb et al., 2009; Gerson-Gurwitz et al., 2016). In contrast, *csr-1* null mutant males are fertile under normal lab culture conditions, but display a progressive loss of fertility over several generations at a stressful temperature (25°C) (Conine et al., 2013). Therefore, we tested whether either isoform was sufficient for viability, or if both were redundantly required for proper germline development and function.

To assess the contribution of *csr-1a* to germline development, we generated additional *csr-1a* mutant alleles, one allele possessing a premature stop codon within the first exon of *csr-1a, csr-1a^[G120*]^*, and a missense allele in the conserved RG repeats of the CSR-1a N-terminus, *csr-1a^[G91R]^* (Fig. S1A, G). We verified that CSR-1b was the only CSR-1 isoform expressed in the *csr-1a^[G120*]^* mutant strain by western blotting protein lysates with a CSR-1 antibody that recognizes both isoforms (Fig. S4A). In parallel, we observed that although CSR-1a^[G91R]^ is expressed in the appropriate tissues (Fig. S4B and data not shown), it is present at lower levels than wild-type CSR-1a. This suggests that *csr-1a^[G91R]^* could behave as a hypomorph (Fig. S4A). Because CSR-1a is not expressed in the hermaphrodite germline or in early embryos, we expected that loss of *csr-1a* would likely have little impact on oocyte development or embryo viability. Furthermore, our GFP::3xFLAG::CSR-1b-expressing strains possess a null mutation in *csr-1a* (*csr-1a(tor159)*) and display wild-type fertility under normal lab culture conditions (Fig. S1B). Indeed, we observed that the brood size and viability of *csr-1a* mutants were comparable to N2 wild-type worms at 20°C (Fig. 2A). At the stressful temperature of 25°C, we observed a slight, but reproducible decrease in brood size in both *csr-1a* mutants relative to wild-type worms. This subtle phenotype at 25°C led us to pursue the role of *csr-1a* at 25°C further, so we cultured the *csr-1a(tor159)* null mutant at 20°C for >25 generations, then shifted it to 25°C to perform brood analysis. In parallel, we freshly outcrossed these “late generation” *csr-1a(tor159)* null mutants, shifted them to 25°C and assayed their brood size in parallel. We found that *csr-1a(tor159)* null mutants in culture for many generations displayed a smaller brood size than wild-type worms at 25°C, but that this phenotype was reversed by outcrossing to N2 (Fig. 2B). This phenotype is reminiscent of transgenerational epigenetic inheritance defects displayed by other germline *ago* mutants, including *csr-1(tm892)* null mutant males, which express neither isoform of *csr-1* (Conine et al., 2013). Collectively, these observations lead us to conclude that under normal culture conditions, CSR-1a is not required for embryonic viability. Instead *csr-1b* plays an essential role during oocyte development and in the early embryo. However, under stressful conditions, such as elevated temperature, *csr-1a* may play a role in fertility that is likely linked to a role in sperm development and differentiation (see below).

**Figure 2.**
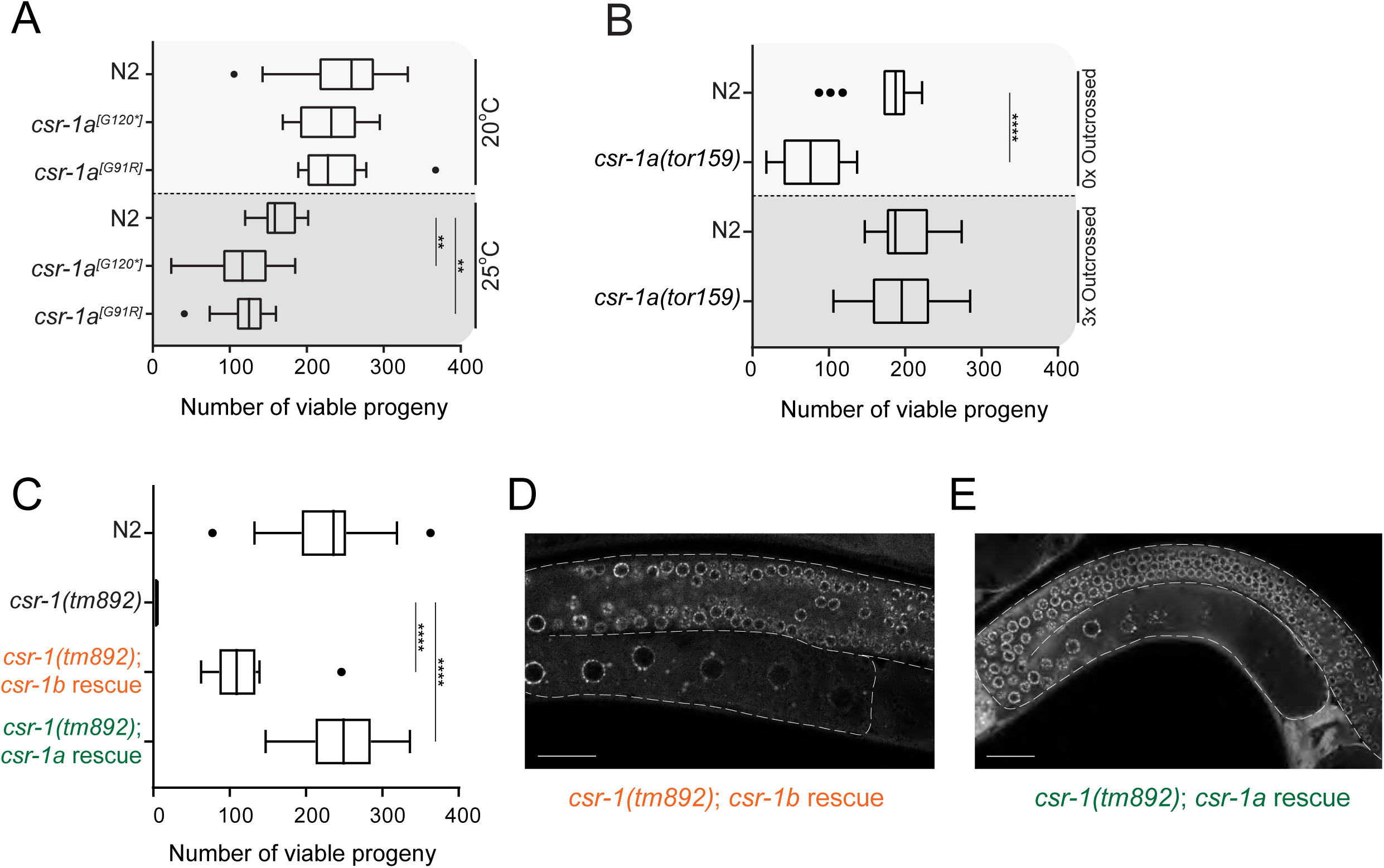
CSR-1b is the essential CSR-1 isoform. **A)** Box plot of viable brood counts for *csr-1a* mutants reared at 20°C and 25°C. Median is represented, with Tukey Whiskers. ** indicates significance of *p*<0.01 and it was determined with a one-way ANOVA (alpha=0.05) with Tukey’s multiple comparison test. Differences are not significant unless marked. N for each genotype and temperature is 14 or more individual P_0_ hermaphrodites. **B)** Box plot of viable brood counts of *csr-1a(tor159), gfp::3xflag::csr-1b* animals reared at 25°C. 0x Outcrossed worms were reared for >25 generations at 20°C before shifting to 25°C for brood assessment. 3x Outcrossed worms were crossed to wild-type animals before shifting to 25°C for brood assessment. Median is represented, with Tukey Whiskers. **** indicates significance of *p*<0.0001 and it was determined with a one-way ANOVA (alpha=0.05) with Tukey’s multiple comparison test. Differences are not significant unless marked. N for each genotype is 16 or more individual P_0_ hermaphrodites. **C)** Box plot of viable brood counts for the *csr-1(tm892)* null mutant rescued with *csr-1b* and *csr-1a* transgenes expressed under control of the *rpl-28* promoter and reared at 20°C. Median is represented, with Tukey whiskers. **** indicates significance of *p*<0.0001 and it was determined with a one-way ANOVA (alpha=0.05) with Tukey’s multiple comparison test. Differences are not significant unless marked. N for each genotype is 13 or more individual P_0_ hermaphrodites. **D)** and **E)** Fluorescence micrographs showing germline expression of GFP::CSR-1 in animals rescued with each *csr-1* transgene. Scale bar, 25μm.

Intending to verify these results with an independent line of investigation, we constructed a set of single copy genomically-integrated transgenes, in which the coding sequence of each *csr-1* isoform was tagged with *gfp* and was expressed using a constitutive *rpl-28* promoter and *csr-1* 3′UTR in a *csr-1(tm892)* null mutant background (Fig. S1A, Fig. 2D-E, Fig. S4C). Note that in these transgenic strains, *csr-1a* was *gfp*-tagged internally, while *csr-1b* was *gfp*-tagged at the N-terminus, because the *gfp* sequence was inserted at the same position in both isoforms. When we constitutively expressed GFP::CSR-1b in the *csr-1(tm892)* null mutant, we found these worms could be maintained as a viable strain, indicating complementation (Fig. 2C). We next measured the extent to which expression of GFP::CSR-1b rescued fertility by performing total brood counts versus viable brood counts. In contrast to *csr-1(tm892)*, which produces very few embryos, this strain produced 73.1% the total brood of wild-type worms (data not shown). However, despite this robust brood size, we found that many of the embryos were not viable (Fig. 2C; *csr-1b* rescue viability was 47.4% of wild-type worms), and died late in embryogenesis as malformed larvae (Fig. S4D). This embryonic arrest occurs later in development than embryos produced by *csr-1(tm892)* homozygous hermaphrodites, suggesting that the phenotype could be due to different causes, including the developmental timing and cell types in which CSR-1b was expressed or the level to which it was expressed.

In a similar set of experiments, we expressed GFP::CSR-1a from the *rpl-28* promoter in the *csr-1(tm892)* null mutant. We performed western blots to verify that only GFP::CSR-1a was expressed from this transgene and that there was no CSR-1b protein present in these strains (Fig. S4C). By using this constitutive promoter, CSR-1a is expressed in many tissues and at developmental time points where and when it would not normally be expressed, including in the oogenic germline and in early embryos (Fig. 2E). Remarkably, this strain was viable, and displayed a viable brood size comparable to wild type (Fig. 2C), indicating that GFP::CSR-1a can compensate for loss of *csr-1b* when ectopically expressed in the oogenic germline and in embryos. Overall, this result was unexpected, and suggests that CSR-1a can execute the molecular functions of CSR-1b, but has differentiated itself from CSR-1b based on its expression profile.

### CSR-1a and CSR-1b associate with distinct pools of sRNAs

To understand whether each CSR-1 isoform has distinct or overlapping target transcripts and roles in gene regulation, we set out to determine the pools of sRNAs bound by each isoform. Past studies have shown that CSR-1 associates with 22G-RNAs, and 22G-RNAs display perfect complementarity to target transcripts (Gu et al., 2009; Claycomb et al., 2009; Conine et al., 2013). Thus, by determining the sRNA complements associated with each CSR-1 isoform, we can computationally identify gene targets. In these experiments, we used GFP antibodies to perform immunoprecipitation (IP) of GFP::3xFLAG::CSR-1 (for CSR-1a, CSR-1b and CSR-1) in synchronized L4 worm populations (Fig. S5). We chose L4 worms because both isoforms are robustly expressed at this stage. For CSR-1b we expected that sRNA populations would reflect germline activities during spermatogenesis. For CSR-1a, we anticipated that we would recover both somatic and germline (spermatogenesis) sRNA populations, due to its expression profile. In parallel, we cloned and sequenced total sRNA samples from the same samples (Input). Two replicates were performed for each condition (Supp. Table S1-2).

We focused our efforts on the CSR-1a and CSR-1b IPs, and first examined the size and first nucleotide distribution in the pool of genome-mapping sRNAs from the Input and IP samples. As anticipated, we observed a large proportion of 22G-RNA reads present in each CSR-1 IP (Fig. 3A, Supp Table S1-2). However, we were surprised to also observe an additional fraction of 26G-RNAs associated with CSR-1a that were not present in the CSR-1b IP (Fig. 3A). 26G-RNAs are present in low proportions in Input samples, and have so far been found in association with ALG-3, ALG-4 and ERGO-1 (Conine et al., 2010; Vasale et al., 2010)

**Figure 3.**
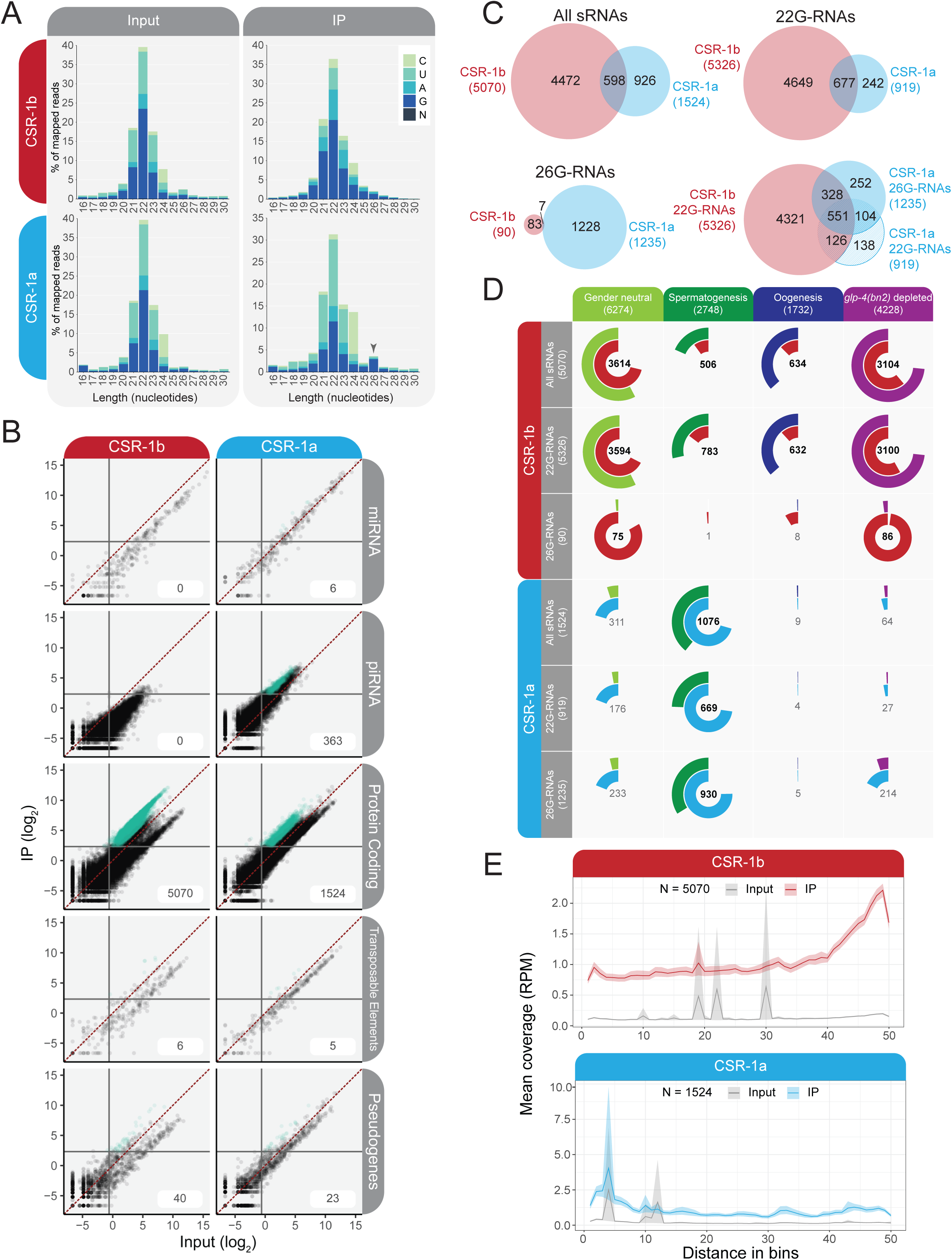
CSR-1b and CSR-1a associate with different pools of sRNAs. **A)** Bar graph of first nucleotide and size distribution of normalized sRNA reads from Input and IP samples sequenced from GFP::3xFLAG::CSR-1b and GFP::3xFLAG::CSR-1a. **B)** Scatter plot of sRNA data showing enrichment in IP relative to Input. Teal dots represent transcripts/genes which have a minimum of 5RPM in IP samples and are 2-fold or greater enriched relative to Input in both replicates. **C)** Venn Diagram of overlap between the enriched sRNAs antisense to protein coding genes from GFP::3xFLAG::CSR-1a and GFP::3xFLAG::CSR-1b IPs. All sRNAs = no size restriction of sRNAs, 22G-RNAs = sRNAs in the range of 21-23nt with no first nucleotide bias, 26G-RNAs = sRNAs in the range of 25-27nt, with no first nucleotide bias. **D)** Venn-pie diagrams comparing CSR-1 enriched sRNAs antisense to protein coding genes (inner circle, red or blue) with gene sets that define spermatogenesis-enriched transcripts, oogenesis-enriched transcripts, and gender neutral transcripts (enriched in both spermatogenesis and oogenesis) (Ortiz et al., 2014), and germline expressed sRNAs (as determined by genes that are 2 fold or more depleted of sRNAs in *glp-4(bn2)* mutants) (Gu et al., 2009) (outer circle, various colors). All sRNAs = no size restriction of sRNAs, 22G-sRNAs = sRNAs in the range of 21-23nt with no first nucleotide bias, 26G-sRNAs = sRNAs in the range of 25-27nt with no first nucleotide bias. The size of each color-coded circle represents the proportion of each sample that is shared between the two sets. Numbers in the center of the circle indicate the number of genes that overlap between the two data sets. Numbers in bold are statistically significant as calculated by Fisher’s Exact Test, *p*<0.001. **E)** Metagene plot of the distribution of enriched sRNAs antisense to protein coding genes along the gene body for the All sRNAs group. N = Number of genes. CSR-1a = GFP::3xFLAG::CSR-1a and CSR-1b = GFP::3xFLAG::CSR-1b.

The observation that CSR-1a associates with 26G-RNAs was unexpected, thus we sought to assess the specificity of this interaction. First, we performed CSR-1a IP-sRNA sequencing using an antibody specific for CSR-1a (Fig. S6). Similar to the GFP::3xFLAG::CSR-1a IP, we observed both 22G-RNAs and 26G-RNAs in this sample (Fig. S6A, and see below). We also assessed whether the 26G-RNAs present in the CSR-1a IP could be due to an association with the 26G-RNA binding AGOs ALG-3 and −4 by performing CSR-1a IPs using the CSR-1 antibody that recognizes both isoforms, and the CSR-1a specific antibody in a GFP::3xFLAG::ALG-3 strain (Fig. S6F). Over multiple experiments, we were unable to observe a physical interaction. Consistent with this result, recently published CSR-1 proteomic data also failed to detect an interaction between either CSR-1 isoform and ALG-3 or ALG-4 (Barucci et al., 2020). [Note: we were unable to assess an interaction with ALG-4 at this point because of research stoppage due to COVID-19.] These data suggest that CSR-1a associates with both 26G-RNAs and 22G-RNAs.

We determined which reads were enriched in each GFP::3xFLAG::CSR-1 IP relative to Input (with a minimum of 5 reads per million (RPM) and a 2-fold or greater increase in IP versus Input) to define a set of target transcripts (genes for which antisense sRNAs are enriched in the IP) for each isoform (Fig. 3B, Supp. Table S2). Consistent with previous studies of CSR-1 (Claycomb et al., 2009; Conine et al., 2013), protein coding genes are the major targets of both isoforms (CSR-1a: 1524 genes; CSR-b: 5070 genes) (Fig. 3B). Both isoforms also target a small fraction of the transposable element complement of *C. elegans* (CSR-1a: 5/268 TE detected in our samples; CSR-b: 6/284 TE detected in our samples) and pseudogenes (CSR-1a: 23/1327 pseudogenes detected in our samples; CSR1-b: 40/1479 pseudogenes detected in our samples) (Fig. 3B). CSR-1a also enriches for a very small number of miRNAs and piRNAs (6/328 miRNAs detected in our samples and 363/12243 piRNAs detected in our samples). Consistent with these results, the CSR-1a antibody-IP sequencing enriched for sRNAs predominantly targeting protein coding genes (4182 genes) as well as a small number of transposable elements (14/275), pseudogenes (92/1169), miRNAs (19/295), and piRNAs (163/9352) (Fig. S6B).

We next asked whether sRNAs associated with each CSR-1 isoform could regulate distinct sets of target genes. Different classes of sRNAs may mediate distinct gene regulatory functions. Therefore, in addition to considering the complete set of sRNA targets described above, we also separately considered sRNAs associated with each CSR-1 isoform within two classes: 22G-RNAs (21-23nt in length, without a 5′ nucleotide bias), and 26G-RNAs (25-27nt in length, without a 5′ nucleotide bias), and used enrichment of these reads to identify targets of each isoform and sRNA group. We note that the numbers of targets of 22G-RNAs and 26G-RNAs do not add up to the number of targets for each CSR-1 isoform when the total pool of sRNAs is considered. This is because some genes are enriched for 22G-RNAs but not 26G-RNAs, some are enriched for 26G-RNAs but not 22G-RNAs, and some genes are enriched for both. From here, we examined the extent that targets overlapped between CSR-1a and CSR-1b. Considering the total pool of sRNAs, we identified 598 genes that are the targets of both CSR-1 isoforms (Fig 3C, Supp. Table S2). We found that 677 genes are the targets of 22G-RNAs bound by both CSR-1 isoforms (Fig 3C). However, as very few 26G-RNAs associate with CSR-1b, only 7 genes are the targets of 26G-RNAs bound by both isoforms (Fig 3C). Rather, the majority of CSR-1a 26G-RNA targets overlap with CSR-1b 22G-RNA targets (879/1235 genes) and/or CSR-1a 22G-RNA targets (655/1235 genes), indicating that CSR-1a 26G-RNAs could function upstream of CSR-1a or CSR-1b 22G-RNAs (Fig. 3C).

The endogenous CSR-1a antibody IP-sRNA sequencing data identified 779 26G-RNA targets, of which almost all (732) were also detected as 26G-RNA targets of the GFP CSR-1a IP (Fig. S6C, Supp. Table S2). We identified more 22G-RNAs in this sample than in the GFP IP, the majority of which are also found in CSR-1b or CSR-1 (total) IPs, indicating that there could be more extensive overlap between the 22G-RNA targets of the two isoforms than detected by the GFP CSR-1a IP (Fig S6D). Overall, these data support a model in which CSR-1a associates with a pool of 26G-RNAs as well as a set of 22G-RNAs that primarily target protein coding genes. Some of these genes are also targeted by CSR-1b 22G-RNA complexes. In addition, CSR-1a and CSR-1b each bind to a large repertoire of small RNAs that are complementary to isoform-specific targets.

### CSR-1a and CSR-1b target different sets of germline-expressed protein coding genes

The distinct targets of CSR-1a and CSR-1b may reflect their distinct expression profiles. To better understand this, we categorized the protein coding gene sRNA targets of each CSR-1 isoform based on existing datasets that designate sperm, gender neutral, and oocyte enriched transcripts (Ortiz et al., 2014), and that examined which sRNAs are expressed in the germline by measuring sRNA depletion in a *glp-4(bn2)* mutant that does not develop a germline at restrictive temperatures (Gu et al., 2009) (Fig. 3D, Supp. Table S2-S3). Consistent with the expression of CSR-1a during spermatogenesis, CSR-1a targets are highly enriched in spermatogenesis genes (1076/1524 all sRNA targets), and are almost devoid of genes expressed during oogenesis (9/1524 all sRNA targets) (Fig. 3D). CSR-1b targets are highly enriched in “gender neutral” genes that are expressed during both spermatogenesis and oogenesis (3614/5070 all sRNA targets). CSR-1b targets also encompass a large proportion of genes that are heavily depleted of sRNAs in mutants possessing no germline (*glp-4(bn2)*) (3104/5070 all sRNA targets), indicating germline-specific expression of these genes. Oogenesis enriched genes are an abundant CSR-1b target group (634/5070 all sRNA targets), and spermatogenesis enriched genes are comparably targeted (506/5070 all sRNA targets) by CSR-1b (Fig. 3D). This is consistent with expression of CSR-1b throughout the germline, and may suggest specialized roles in spermatogenesis and oogenesis. Notably, very few CSR-1a targets were depleted in *glp-4(bn2)* samples (64/1524 all sRNA targets). This indicates that although they are spermatogenesis-enriched, many targets of CSR-1a are expressed in somatic cells, and is consistent with the expression of CSR-1a in somatic tissues and during spermatogenesis. The endogenous CSR-1a antibody IP displayed a similar profile of enrichment for spermatogenesis genes (Fig. S6E) (1413/4182 all sRNA targets, 1306/3815 22G-RNA targets, and 592/779 26G-RNA targets). The additional 22G-RNA targets observed in this experiment allowed for the identification of a greater number of gender neutral genes among CSR-1a targets (2117/4182 gender neutral genes for all sRNA targets; 1985/3815 for 22G-RNA targets) and a greater number of *glp-4(bn2)* depleted genes (1439/4182 for all sRNA targets; 1436/3815 for 22G-RNA targets). We conclude that CSR-1a largely targets spermatogenesis-enriched genes, although these targets may also be expressed (and targeted) in the soma. In contrast, CSR-1b targets a distinct set of gender neutral, germline-specific genes.

We next asked if there are any differences in the distribution of sRNAs along the groups of target genes for each CSR-1 isoform. To assess this, we performed metagene analysis, examining the density of sRNAs along the group of CSR-1a vs. CSR-1b protein coding gene targets, after normalizing for gene length (Fig. 3E, Fig. S7A). CSR-1b-associated sRNAs displayed a pattern consistent with previous reports, in which sRNAs were enriched at the 3′ end of the target transcripts. However, we observed a distinct pattern of sRNAs associated with CSR-1a, in which sRNAs were enriched at the 5′ end of the target genes. This result is reminiscent of the pattern of sRNAs associated with the other 26G-RNA binding spermatogenesis AGOs, ALG-3 and ALG-4. We therefore were curious if this pattern was present for either the 22G-RNAs or the 26G-RNAs associated with CSR-1a. After dividing the sRNA reads into these two groups we found that the 5′ bias in sRNA reads was present in both. Hence this is another feature of CSR-1a that distinguishes it from CSR-1b, and is not related only to CSR-1a’s association with 26G-RNAs.

Finally, we examined a unique set of protein coding transcripts, the histone genes. Replication dependent histone mRNAs are generally not poly-adenylated, but possess a 3′ stem loop that must be cleaved for mRNA maturation and translation (Keall et al., 2007). CSR-1 has been implicated in histone mRNA processing in worms. In this mechanism, sRNAs guide CSR-1 to the stem loop cleavage site, where CSR-1 is proposed to act as the endonuclease (Avgousti et al., 2012). There are approximately 70 histone genes detectable in the *C. elegans* genome and we found that 50% of the histone genes are targeted by CSR-1a, CSR-1b, or both isoforms (Fig. S7B, Supp. Table S2). Each isoform targets distinct subsets of histones predominantly by 22G-RNAs. CSR-1a targets H2B, H3.2, H3.3 and H4, while CSR-1b targets predominantly H2A (including important variants like H2AX, which is involved in DNA damage). H4 is a prominent target of both isoforms. Because both CSR-1 isoforms possess the same catalytic tetrad within the PIWI domain, it is plausible that both isoforms could process the 3′ stem loop of different classes of histone genes in different tissues during development.

### CSR-1b and CSR-1a differentially intersect with the WAGO-4 pathway

Recent studies have demonstrated that the Worm AGO, WAGO-4 binds to a pool of 22G-RNAs that target nearly all of the previously defined CSR-1 protein coding gene targets (Xu et al., 2018). WAGO-4 is expressed constitutively in the germline, where it localizes to germ granules, including Z granules in adults and P granules in early embryos. WAGO-4 has been implicated in transgenerational epigenetic inheritance of exogenous RNAi and its loss leads to a Mortal germline (Mrt) phenotype, in which fertility declines over successive generations at 25°C (Xu et al., 2018; Wan et al., 2018; Ishidate et al., 2018). Because of these data linking the CSR-1 and WAGO-4 pathways, we aimed to delineate the relationship between each CSR-1 isoform and WAGO-4.

WAGO-4 associated sRNAs and targets have been identified previously (Xu et al., 2018). We also developed our own GFP::3xFLAG::WAGO-4 associated sRNA data sets, and compared these data to the targets of CSR-1a and CSR-1b here. Our WAGO-4 IPs identified 5021 WAGO-4 all sRNA targets and show strong overlap with existing data sets (Xu et al., 2018) (81.4% overlap, 4061/5021 WAGO-4 targets defined by our experiments overlap with the 4748 previously defined WAGO-4 targets). Of these 5021 WAGO-4 targets, 4999 are targeted by 22G-RNAs. Here, we found that approximately 90% of the CSR-1b (4539/5070) and WAGO-4 (4539/5021) targets of all sRNAs overlap (Fig. S7C, Supp. Table S4). In contrast, only 13% (198/1524) of the CSR-1a all sRNA targets and 4% (198/5021) of the WAGO-4 all sRNA targets overlap (Fig. S7C). Among the 198 target genes shared between CSR-1a and WAGO-4, 186 of these genes are also targeted by CSR-1b (Fig. S7C). We next considered the separate pools of CSR-1a 22G-RNAs and 26G-RNAs, and found that more 26G-RNA targets overlapped with WAGO-4 targets; almost 23% (281/1235) of CSR-1a 26G-RNA target genes overlapped with WAGO-4 target genes vs. approximately 7% (95/1235) of CSR-1a 22G-RNA targets (Fig. S7C). This indicates that CSR-1a is likely to act in an upstream or primary capacity to WAGO-4, as it may with CSR-1b. Collectively, these data indicate that CSR-1b and WAGO-4 coordinately regulate target transcripts in the germline via 22G-RNAs, but that CSR-1a plays a distinct role with its 22G-RNAs and may operate in a primary capacity to WAGO-4 via 26G-RNAs.

To further understand the relationship between CSR-1 isoforms and WAGO-4, we examined the uridylation status of sRNAs associated with each CSR-1 isoform. CSR-1 22G-RNAs are known to be uridylated at their 3′ ends by terminal uridyl transferase, CDE-1, and this modification was originally thought to facilitate turnover of CSR-1-associated 22G-RNAs (van Wolfswinkel et al., 2009; Claycomb et al., 2009). Conversely, WAGO-4 favors an association with the uridylated forms of these 22G-RNAs, hence uridylation of the 22G-RNAs may control the balance in association between CSR-1 and WAGO-4 (Xu et al., 2018). We examined the 22G-RNAs and 26G-RNAs associated with each CSR-1 isoform for evidence of untemplated U addition at the 3′ end (Fig. S7D, Supp. Table S2). We found that CSR-1b 22G-RNAs were consistently U-tailed (18% of 22G-RNA reads in IP samples had one or more U added vs. 10% in Input samples). Although extremely low in number, CSR-1b associated 26G-RNAs also displayed a similar degree of U-tailing (16% of reads in IP samples vs. 11% in Input samples). In contrast, CSR-1a 22G-RNAs displayed a more modest degree of U-tailing (12% of 22G-RNA reads in IP samples vs. 8% in Input samples), and 26G-RNAs showed no evidence of U-tailing (9% of 26G-RNAs in IP samples vs. 8% in Input samples). These differences in the U-tailing of CSR-1b-versus CSR-1a-associated sRNAs uncover another important distinction between the CSR-1 isoforms, and suggest that CSR-1a-associated sRNAs are regulated differently than those in complex with CSR-1b.

Collectively, our results support a model in which the CSR-1b and WAGO-4 22G-RNA pathways coordinately regulate mostly gender neutral genes that are constitutively expressed during both spermatogenesis and oogenesis. The CSR-1a pathway predominantly regulates a separate set of spermatogenesis-enriched transcripts via 22G-RNAs and 26G-RNAs, and has the potential to act upstream of (or in parallel to) CSR-1b and WAGO-4 via its 26G-RNAs for a subset of shared target genes.

### CSR-1a is required for spermatogenesis under temperature stress

In a related but separate study, we evaluated the expression patterns and sRNA binding partners of every *C. elegans* AGO (Seroussi, in preparation). This study identified another novel WAGO with a spermatogenesis specific expression profile, WAGO-10 (Fig. 4A). WAGO-10 also associates with a subset of 26G-RNAs like the other spermatogenesis-specific AGOs, ALG-3, ALG-4 and CSR-1a (Seroussi, in preparation). We compared the 26G-RNA targets of each of these spermatogenesis AGOs to understand whether they target distinct or overlapping groups of genes during spermatogenesis. By unsupervised clustering analysis, we found that WAGO-10-associated 26G-RNAs target a relatively small set of genes (447) that overlapped entirely or almost entirely with ALG-3 (447/447), ALG-4 (446/447), and CSR-1a (442/447) 26G-RNA target gene sets (Fig. 4B, Fig. S8A-B, Supp. Table S4). This analysis defines a core set of 442 genes targeted by all spermatogenesis AGOs via 26G-RNAs (irrespective of downstream 22G-RNA binding AGOs). As expected, these 442 targets are enriched in spermatogenesis genes (Fig. S8C) and encompass GO molecular functions that overwhelmingly encompass phosphatase activity, along with cellular compartment enrichment of pseudopodium, cytoskeleton, and intracellular non-membrane bounded organelle (Supp. Table S5), all features known to be essential for differentiating and activating fertile spermatozoa.

**Figure 4.**
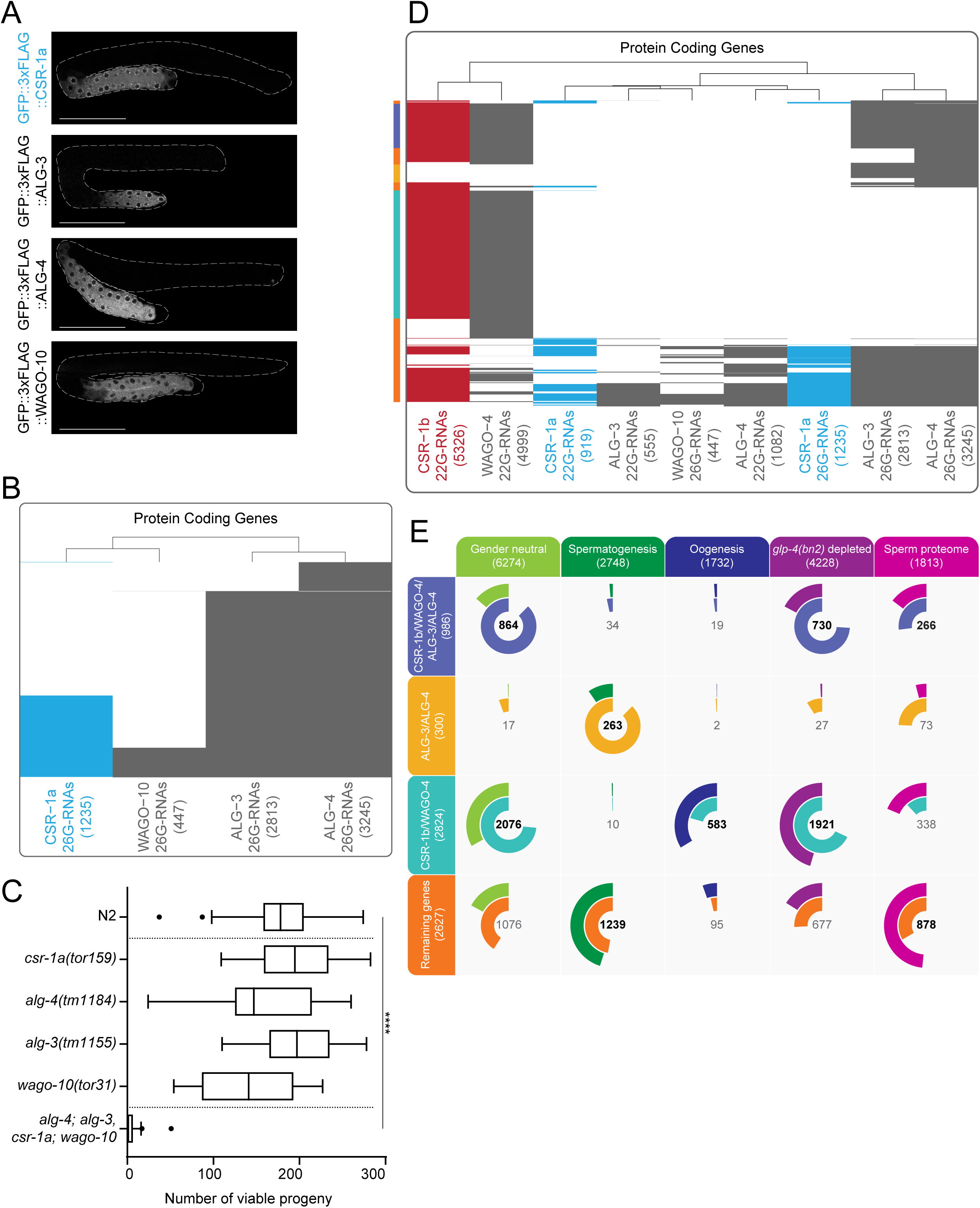
CSR-1a integrates into spermatogenesis 26G-RNA pathways. **A)** Fluorescence micrographs showing L4 germlines of GFP::3xFLAG tagged CSR-1a, ALG-3, ALG-4, and WAGO-10. Scale bar, 50μm. **B)** Clustering diagram of genes targeted by 26G-RNAs (26G-sRNAs = sRNAs in the range of 25-27nt with no first nucleotide bias) enriched in IPs of each spermatogenesis AGO. **C)** Box plot of median viable brood size counts for spermatogenesis *ago* mutants (with Tukey whiskers). Single mutants were outcrossed before brood analysis and the quadruple *ago* mutant was outcrossed in the course of generating the strain. Significance was determined with a one-way ANOVA (alpha=0.05) with Tukey’s multiple comparison test. **** indicates significance of *p*<0.0001, and refers to all pairwise contrasts with the quadruple *ago* mutant. N for each genotype is 12 or more individual P_0_ hermaphrodites. **D)** Clustering diagram of genes targeted by 22G-RNA binding AGOs and 26G-RNA binding spermatogenesis AGOs. This analysis delineates four major groups of genes that are differentially targeted by AGOs and examined in E). Note that the colors along the left side match to gene sets examined in E). **E)** Venn-pie diagrams comparing groups of genes targeted by different sRNAs and AGOs (inner circle) with gene sets that define spermatogenesis-enriched transcripts, oogenesis-enriched transcripts, and gender neutral transcripts (enriched in both spermatogenesis and oogenesis) (Ortiz et al., 2014), germline expressed sRNAs (as determined by genes that are 2 fold or more depleted of sRNAs in *glp-4(bn2)* mutants) (Gu et al., 2009), and sperm proteome-enriched genes (outer circle). The size of each color-coded circle represents the proportion of each sample that is shared between the two sets. Numbers in the center of the circle indicate the number of genes that overlap between the two data sets. Numbers in bold are statistically significant as calculated by Fisher’s Exact Test, *p*<0.001.

The extensive overlap in target genes suggested that these four AGOs may function redundantly. Therefore, we built a quadruple null mutant for the 26G-RNA binding spermatogenesis AGOs, and tested male fertility. Double mutants of *alg-3; alg-4* and *csr-1* null mutants have previously been shown to exhibit a progressive loss of male fertility over several generations when cultured at the stressful temperature of 25°C (Conine et al., 2010; Conine et al., 2013). We also noticed that our *csr-1a(tor159)* allele showed a decrease in fertility at 25°C after long-term culture at 20°C (Fig. 2B). Thus, we outcrossed each single spermatogenesis *ago* mutant and compared the brood size of each freshly outcrossed single mutant to the quadruple spermatogenesis *ago* mutant in the first generation experiencing the temperature shift. For each of the outcrossed single mutants, the brood size was comparable to wild-type worms (Fig. 4C). In stark contrast, the quadruple *ago* mutant was nearly sterile in the first generation experiencing temperature shift (Fig. 4C). To determine whether the fertility defect was due to non-functional sperm, we crossed the quadruple spermatogenesis *ago* mutant to wild-type males, and found that this rescued the fertility defect (Fig. S8D). Collectively, these data indicate that 26G-RNA binding spermatogenesis AGOs function redundantly to regulate a core set of spermatogenesis genes necessary for fertility.

### An interconnected network of AGOs regulate germline transcripts during spermatogenesis

The ALG-3/-4 26G-RNA pathway was previously described as acting in an upstream or primary manner to the CSR-1 and WAGO 22G-RNA pathways (Conine et al., 2010; Conine et al., 2013). A survey of our sRNA data for all AGOs (Seroussi, in preparation) indicated three critical features about these pathways that we explored further. First, ALG-3 26G-RNA target genes almost entirely coincide with ALG-4 26G-RNA targets (2807/2813 ALG-3 targets overlap with the set of 3245 ALG-4 targets) (Fig. S8A). Second, like CSR-1a, we observed that ALG-3 and ALG-4 bind to both 26G-RNAs and 22G-RNAs. ALG-3 and ALG-4 IPs enrich for 22G-RNAs that target 555 and 1082 genes, respectively. Third, CSR-1b and WAGO-4 were the two 22G-RNA binding AGOs expressed during spermatogenesis and possessing the most extensive overlap with the targets of ALG-3 and ALG-4 26G-RNAs. This suggests that CSR-1b and WAGO-4 may be the predominant secondary 22G-RNA associated AGOs that act downstream of the spermatogenesis AGOs that associate with 26G-RNAs (Fig. 4D, Supp. Table S4).

To better delineate the relationships between 22G-RNA binding AGOs and 26G-RNA binding AGOs, we performed an unsupervised clustering analysis to identify subsets of genes that were coordinately targeted by multiple AGOs (Fig. 4C-D, Supp. Table S4-S5). Several distinct groups of genes emerged from this analysis. First, we observed a set of 986 genes that are targeted by ALG-3 and ALG-4 26G-RNAs, along with CSR-1b and WAGO-4 22G-RNAs (Fig. 4C-D, Supp. Table S4). These genes fell into the category of gender neutral genes that are depleted in *glp-4(bn2)* worms (Gu et al., 2009) (Fig. 4D, Supp. Table S4). Gene Ontology (GO) analysis (Raudvere et al., 2019) revealed RNA binding and protein binding as the most enriched molecular functions for this group (Supp. Table S5). Second, we identified a set of 300 genes that are only targeted by ALG-3 and ALG-4 26G-RNAs (Fig. 4C-D, Supp. Table S4). These targets are enriched in spermatogenesis genes, and the most enriched GO term was protein tyrosine phosphatase activity (Fig. 4D, Supp. Table S4-S5). Third, we found the largest group of 2824 genes to be targeted by only CSR-1b and WAGO-4 22G-RNAs (Fig. 4C-D, Supp. Table S4). These genes encompass gender neutral genes, along with a subset of oogenesis genes, and are depleted in *glp-4(bn2)* worms (Gu et al., 2009) (Fig. 4D, Supp. Table S4). GO analysis (Raudvere et al., 2019) showed a strong enrichment of RNA and nucleic acid binding activity, RNA processing, DNA repair, and other gene expression related processes for this set of genes (Supp. Table S4-S5). Finally, the rest of the spermatogenesis AGO target genes (2627 genes) were targeted by single or multiple AGOs and types of sRNAs, including the ALG-3/-4 and CSR-1a 22G-and 26G-RNAs in conjunction with CSR-1b and WAGO-4 22G-RNAs (Fig. 4C-D, Supp. Table S4). These targets may be more heterogeneous in nature, yet a theme still emerges. These genes encompass spermatogenesis genes and gender neutral genes in comparable proportion (Fig. 4D, Supp. Table S4). In addition, a subset of these targets encode proteins identified within the sperm proteome (Ma et al., 2014), and are mildly depleted in *glp-4(bn2)* worms (Gu et al., 2009) (Fig. 4D, Supp. Table S4). GO analysis (Raudvere et al., 2019) of this set of genes revealed an enrichment for molecular functions of protein phosphatase activity, kinase activity, and protein metabolism (Supp. Table S5).

We further examined genes involved in several important facets of spermatogenesis and spermiogenesis in greater detail in an attempt to understand how they might be regulated (Supp. Table S4). CSR-1b and WAGO-4 appear to play broad roles in the spermatogenic germline. All Delta/Notch signaling factors involved in maintaining the germline stem cell niche (White-Cooper and Bausek, 2010) are targeted almost exclusively by CSR-1b and WAGO-4 22G-RNAs (*gld-2* is also a target of CSR-1a, ALG-3, and ALG-4 26G-RNAs). The notable exception in this pathway is *lag-2*, the Delta ligand expressed in the distal tip cell, which is not targeted by any sRNA pathways described here. Consistent with this, Notch signaling is a significant term in the GO analysis (Raudvere et al., 2019) above. Similarly, the sex determination pathway (Ellis and Schedl, 2007) is almost exclusively targeted by CSR-1b and WAGO-4 22G-RNAs, as are many chromatin associated DNA and RNA binding proteins expressed in mature spermatozoa (Chu et al., 2006).

Conversely, sperm-specific functions are regulated more heavily by the spermatogenesis AGOs (Supp. Table S4). Genes involved in spermiogenesis—the maturation and activation of spermatids into motile, fertilization-competent spermatozoa (L’Hernault, 2006; Smith, 2014)—are predominantly targeted by ALG-3, ALG-4, and CSR-1a 26G-RNAs, CSR-1a and ALG-4 22G-RNAs, and to a lesser extent CSR-1b 22G-RNAs. The Major Sperm Proteins (MSPs) are encoded by a group of about 30 highly conserved genes (along with about 20 pseudogenes). These small (∼15kDa), basic proteins are expressed late in spermatogenesis, and play roles in sperm structure, motility, and signaling (Smith, 2014). The *msp* genes fall into the core group of 442 genes targeted by all four 26G-RNA associated spermatogenesis AGOs. They are also targeted to a lesser degree by 22G-RNAs associated with these same AGOs, as well as CSR-1b. Notably WAGO-4 does not target any *msp* genes. While these few examples only scratch the surface, they illustrate the regulatory potential and complexity of sRNA pathways during sperm development.

### Gene expression is perturbed in *csr-1a* mutants

Our work attributes the previously described expression pattern and essential functions of CSR-1 to CSR-1b. We therefore focused our efforts on understanding how loss of CSR-1a affects the expression of its sRNA target genes, and in turn fertility. Because *csr-1a* mutants exhibit temperature-dependent fertility defects after long-term culture, we performed mRNA-seq on late generation L4 stage *csr-1a(tor159)* mutants vs. wild-type worms at 20°C and 25°C.

We first compared the *csr-1a* mutants at 25°C vs. 20°C and found that approximately the same number of genes were two-fold or more up-regulated (167) as were two-fold or more down-regulated (175) (Fig. 5A, Supp. Table S6) (Anders and Huber, 2010; Love et al., 2014). When we overlapped the CSR-1a sRNA targets with the mRNA-seq data, we found that the CSR-1a sRNA targeted genes tended to be up-regulated (Fig. 5A, blue dots). This suggests that CSR-1a, in at least some capacity, silences its target transcripts via sRNAs.

**Figure 5.**
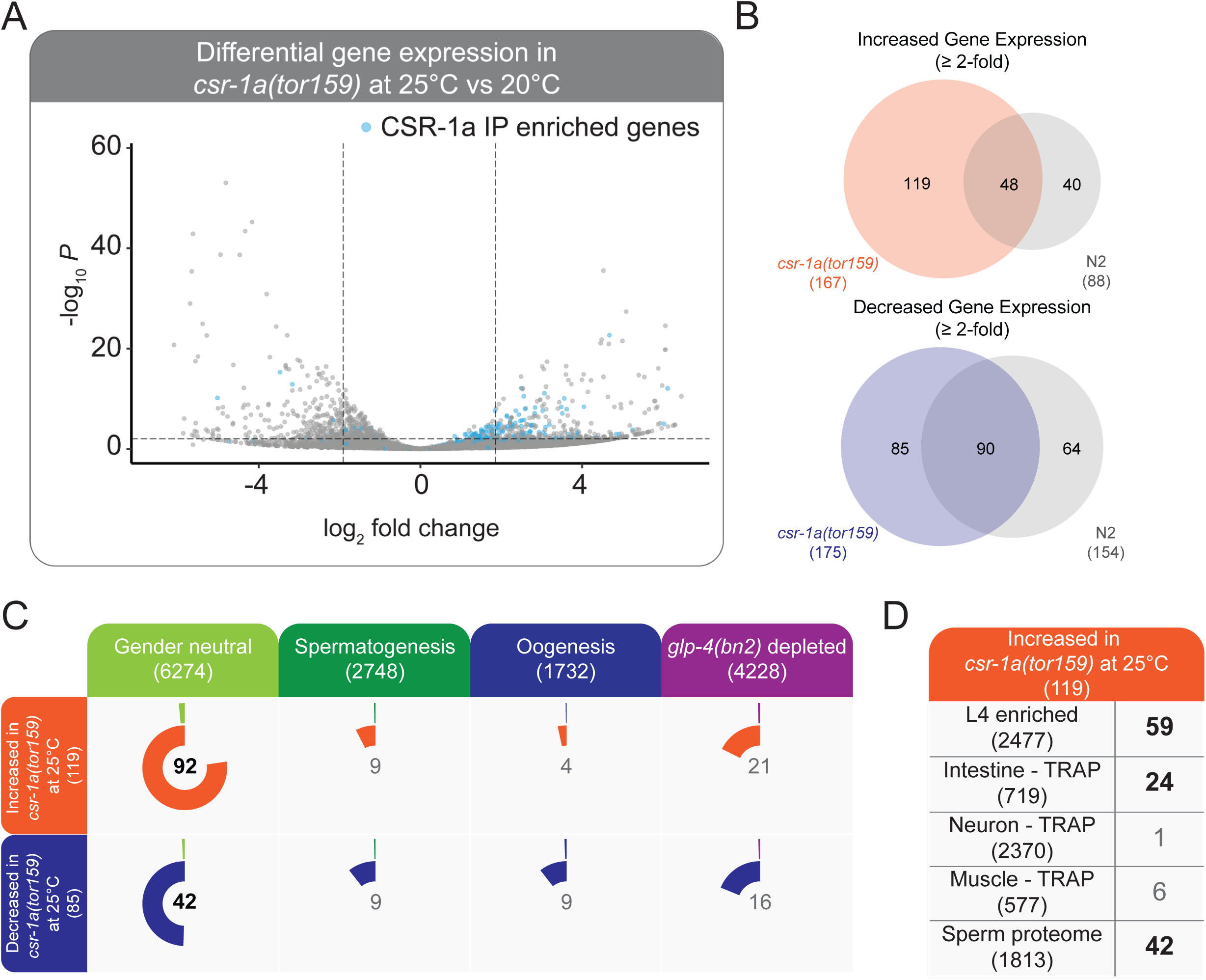
Gene expression analysis in *csr-1a(tor159)*. **A)** Volcano plot of gene expression changes in late generation *csr-1a(tor159), gfp::3xflag::csr-1b* worms reared at 25°C vs. 20°C. Lines denote a *p_adj_* of <0.01 and >2 log_2_-fold change. Blue dots represent genes which are enriched in the GFP::3xFLAG::CSR-1a (denoted as CSR-1a) IP. **B)** Venn Diagrams representing the overlap of genes with expression changes in the *csr-1a(tor159), gfp::3xflag::csr-1b* at 25°C vs 20°C relative to the wild-type (N2) strain at 25°C vs 20°C. We subtracted genes that were altered due to temperature in the N2 strain from our data sets. **C)** Venn-pie diagrams comparing genes that are differentially expressed in *csr-1a(tor159), gfp::3xflag::csr-1b* (inner circle) with gene sets that define spermatogenesis-enriched transcripts, oogenesis-enriched transcripts, and gender neutral transcripts (enriched in both spermatogenesis and oogenesis) (Ortiz et al., 2014), and germline expressed sRNAs (as determined by genes that are 2 fold or more depleted of sRNAs in *glp-4(bn2)* mutants) (Gu et al., 2009) (outer circle). The size of each color-coded circle represents the proportion of each sample that is shared between the two sets. Numbers in the center of the circle indicate the number of genes that overlap between the two data sets. Numbers in bold indicate statistically significant overlap as calculated by Fisher’s Exact Test, *p*<0.001. **D)** Overlap between genes increased in expression in *csr-1a(tor159), gfp::3xflag::csr-1b* versus various data sets that delineate transcripts enriched in L4 hermaphrodites vs. embryos, transcripts engaged with the ribosome in Intestines, Neurons, and Muscles (Gracida et al., 2017), and the sperm proteome genes (Ma et al., 2014). Numbers in bold indicate statistically significant overlap as calculated by Fisher’s Exact Test, *p*<0.001.

To identify genes that changed due to genotype and not because of temperature, we removed the genes that were also differentially expressed in wild-type worms at 25°C vs 20°C from the *csr-1a*-dependent differentially expressed genes (Fig. 5B, Supp. Table S6). This led to final differentially expressed gene sets of 119 up-regulated and 85 down-regulated genes in *csr-a* mutants (Fig. 5B). Of these differentially expressed genes, almost none of the down-regulated genes were CSR-1a sRNA targets (1/85), while a reasonable proportion of the up-regulated genes were CSR-1a sRNA targets (45/119). As a first step to understand the types of genes that were differentially regulated, we overlapped the differentially expressed genes with germline gene expression sets as we had done for our sRNA analyses. We found that for both up-and down-regulated genes, the largest overlap was with gender neutral germline-expressed genes (for up-regulated genes: 92/119 genes; for down-regulated genes: 42/85 genes) (Fig. 5C, Supp. Table S6) (Ortiz et al., 2014).

We went on to examine these sets of differentially expressed genes more carefully, first by performing GO analysis (Raudvere et al., 2019). For the genes with decreased expression, there were no GO terms enriched, and the majority of genes in this set (56/85) only possess WormBase designations rather than gene names reflecting a phenotype, indicating that these are likely to be genes that have not been characterized (Supp. Table S6). Conversely, the genes displaying increased expression showed a weak association with muscle, intestine, and neuronal GO terms (data not shown). To understand which tissues or processes the up-regulated genes were associated with, we examined several additional datasets, including modENCODE mRNA-seq data from each stage of development, Translating Ribosome Affinity Purification (TRAP) data from muscles, neurons, and intestines (Gracida et al., 2017), and sperm proteome data (Ma et al., 2014) (Supp. Table S6). As anticipated based on the stage at which we collected our mRNA samples, 59/119 up-regulated genes were enriched in expression in the L4 stage versus embryos (Fig. 5D, Supp. Table S6). 24/119 up-regulated genes overlapped with genes expressed in the intestine (as determined by TRAP (Gracida et al., 2017)), consistent with intestinal expression of CSR-1a and a potential role for CSR-1a in somatic gene regulation (see below) (Fig. 5D, Supp. Table S6). In contrast, only 6/119 and 1/119 up-regulated genes were present in the muscle and neuronal TRAP data sets (Gracida et al., 2017), respectively (Fig. 5D, Supp. Table S6). Consistent with the expression of CSR-1a during spermatogenesis, 42/119 genes were present in sperm proteome data (Fig. 5D, Supp. Table S6). Importantly, the up-regulated intestinal genes did not overlap with the upregulated sperm proteome genes (Supp. Table S6). Finally, 28 of the transcripts with increased expression encode protein constituents of the ribosome and stood out as the largest single functional group (Supp. Table S6). Of these 28 ribosomal proteins, 23 are enriched in the sperm proteome, and 27 are enriched in mRNA expression during L4 versus embryos (Supp. Table S6). Collectively, these data point to a sRNA dependent role for CSR-1a in silencing different sets of target transcripts in the two major tissues where it is expressed, including a large set of spermatogenesis-expressed ribosomal constituents and intestinal proteins. However, our data do not rule out sRNA independent mechanisms or positive gene regulatory functions.

### CSR-1a plays a role in somatic transgene silencing

In a separate project aimed at understanding the regulation of ubiquitin mediated proteolysis, we serendipitously uncovered a role for CSR-1a in somatic transgene regulation. In these experiments, we generated a genome-integrated repetitive transgene array that drives GFP expression under control of the promoter of the *rpn-2* gene (*rpn-2_p_::gfp*). *rpn-2* encodes the *C. elegans* orthologue of Rpn2/PSMD1, a non-ATPase subunit of the 19S proteasome that is essential for viability of eukaryotic cells (Bard et al., 2018) (Fig. 6A). *rpn-2* is an essential gene in *C. elegans* and is expressed in most or all tissues throughout development (Mounsey et al., 2002; Takahashi et al., 2002). In contrast to the anticipated ubiquitous expression pattern, we found that *rpn-2_p_::gfp* was only weakly expressed in a subset of intestinal cells in otherwise wild-type animals (Fig. 6D). This may be because the promoter fragment used in *rpn-2_p_::gfp* does not encompass all of the regulatory elements that control endogenous *rpn-2* expression. Alternatively, we hypothesized that *rpn-2_p_::gfp* may be subject to the reasonably rare phenomenon of somatic transgene silencing. Notably, transgene silencing is a process that is generally limited to the germline (Grishok et al., 2005; Leyva-Díaz et al., 2017; Lehner et al., 2006; Fischer et al., 2013). The expression of the *rpn-2_p_::gfp* fusion gene was known to be activated in the intestine by proteasome inhibitors or mutations in proteasome subunits, like another proteasomal fusion gene *rpt-3p::gfp* (data not shown and (Lehrbach and Ruvkun, 2016)).

**Figure 6.**
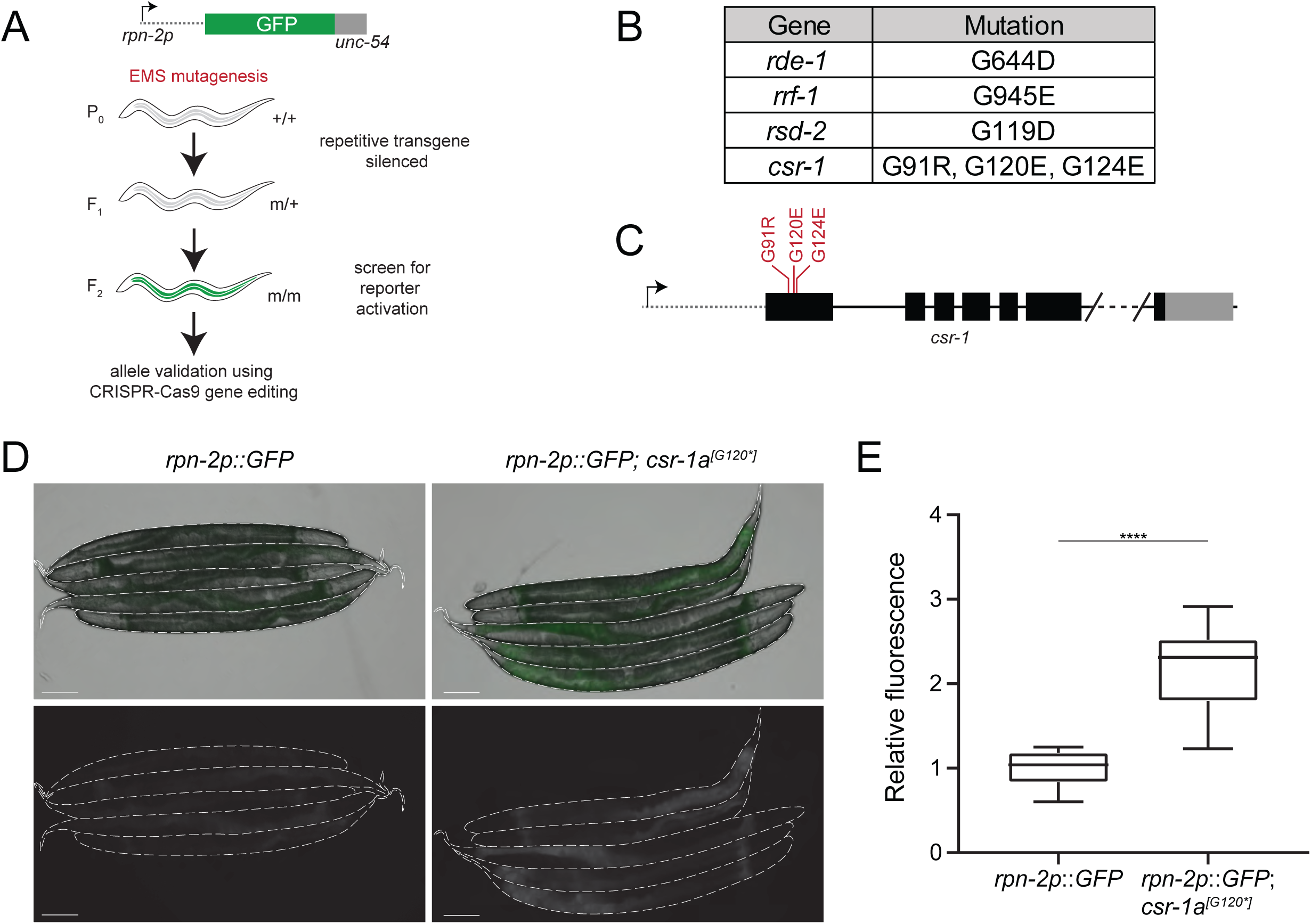
CSR-1a silences a repetitive somatic transgene. **A)** Schematic of the forward genetic screen used to uncover genes that repress *rpn-2p::GFP* expression. **B)** Viable alleles recovered from the forward genetic screen. **C)** Schematic representation of the *csr-1a* alleles recovered from the forward genetic screen. **D)** Fluorescence micrographs of the non-mutagenized reporter strain and the reporter after introduction of *csr-1a^[G120*]^.* Scale bar, 100μm. **E)** Box plot of normalized median relative fluorescence of the reporter strain and the reporter with *csr-1a^[G120*]^* strain (with Tukey whiskers). **** indicates significance of *p*<0.0001 and it was determined with an unpaired *t*-test (alpha=0.05). (*rpn2p::GFP* N=19, *rpn-2p::GFP;csr-1a^[G120*]^* N=16).

To identify additional factors that control expression of the *rpn-2_p_::gfp* transgene and, more generally, regulate the proteasome pathway, we carried out an EMS mutagenesis screen and isolated mutants that show altered patterns of GFP expression (Fig. 6A-D). This screen yielded eleven mutants that show elevated expression of GFP in intestinal cells. We subjected these mutants to whole genome sequencing and have identified the genetic lesion underlying increased GFP expression in seven of the mutants (Fig. 6B-C). Four mutants remain uncharacterized. The alleles we recovered include mutations in the *rrf-1*, *rsd-2* and *rde-1* genes, which encode RNAi pathway factors (Sijen et al., 2001; Tabara et al., 1999; Tijsterman et al., 2004). We confirmed that loss of *rrf-1* and *rde-1* function causes increased expression of the *rpn-2_p_::gfp* transgene using independently derived *rrf-1(pk1417)* and *rde-1(ne300)* alleles (data not shown). RSD-2 acts in concert with the tudor domain protein RSD-6 to promote endogenous and exogenous siRNA-dependent gene silencing (Tijsterman et al., 2004; Zhang et al., 2012; Sakaguchi et al., 2014). Because of the relationship between RSD-2 and RSD-6, we tested an independent allele of *rsd-6(pk3300)* to determine its impact on *rpn-2_p_::gfp* expression, and found that GFP expression is increased in the intestinal cells of *rsd-6(pk3300)* mutant animals (data not shown). These data indicate that the *rpn-2_p_::gfp* transgene is silenced by an siRNA-dependent mechanism in the intestinal cells of wild-type animals that can be relieved by mutations that disable core RNA interference factors such as the AGO RDE-1 or the RRF-1 RNA dependent RNA polymerase.

In addition, we isolated four independent alleles of *csr-1* that all lie within the *csr-1a*-specific first exon (Fig. 6B-C). All of the *csr-1a* alleles recovered cause amino acid substitutions at conserved glycine residues (G91R, two alleles of G120E, G124E) within or nearby RGG/RG repeats of the N-terminus. This specificity and high degree of conservation of these residues across *Caenorhabditis* (Fig. S1G) points to important roles for this motif in the intestine, where *rpn-2_p_::gfp* is expressed. To independently test whether loss of *csr-1a* function causes increased expression of *rpn-2_p_::gfp*, we used CRISPR/Cas9 gene editing to introduce a *csr-1a*-specific stop codon (G120*) to otherwise wild-type *rpn-2_p_::gfp* transgenic animals. We found that *rpn-2_p_::gfp* expression is increased in intestinal cells of animals lacking *csr-1a* (Fig. 6D-E). We conclude that CSR-1a, similar to the RNAi factors identified above, is required for silencing of the repetitive *rpn-2_p_::gfp* transgene in intestinal cells.

## DISCUSSION

Because it is the only essential AGO in *C. elegans*, exhibits several noteworthy developmental and molecular phenotypes, and regulates a large portion of the transcriptome, CSR-1 has been the focus of intense study since its initial characterization. Remarkably, CSR-1 is only conserved in clade III and V nematodes, yet in the only other nematode species in which its function has been explored, *C. briggsae*, it is also essential (Tu et al., 2015). The essential nature of CSR-1 points to important gene regulatory roles in gametogenesis and embryonic development, but until our study, it was unclear which isoforms of CSR-1 were responsible for these varied gene regulatory functions. Using CRISPR-Cas9 genome editing to GFP::3xFLAG tag each CSR-1 isoform, and mutant studies to understand the outcome of loss of each isoform on development, we have uncovered distinct expression patterns, sRNA binding partners, and gene regulatory functions for the two CSR-1 isoforms (Fig. 7).

**Figure 7.**
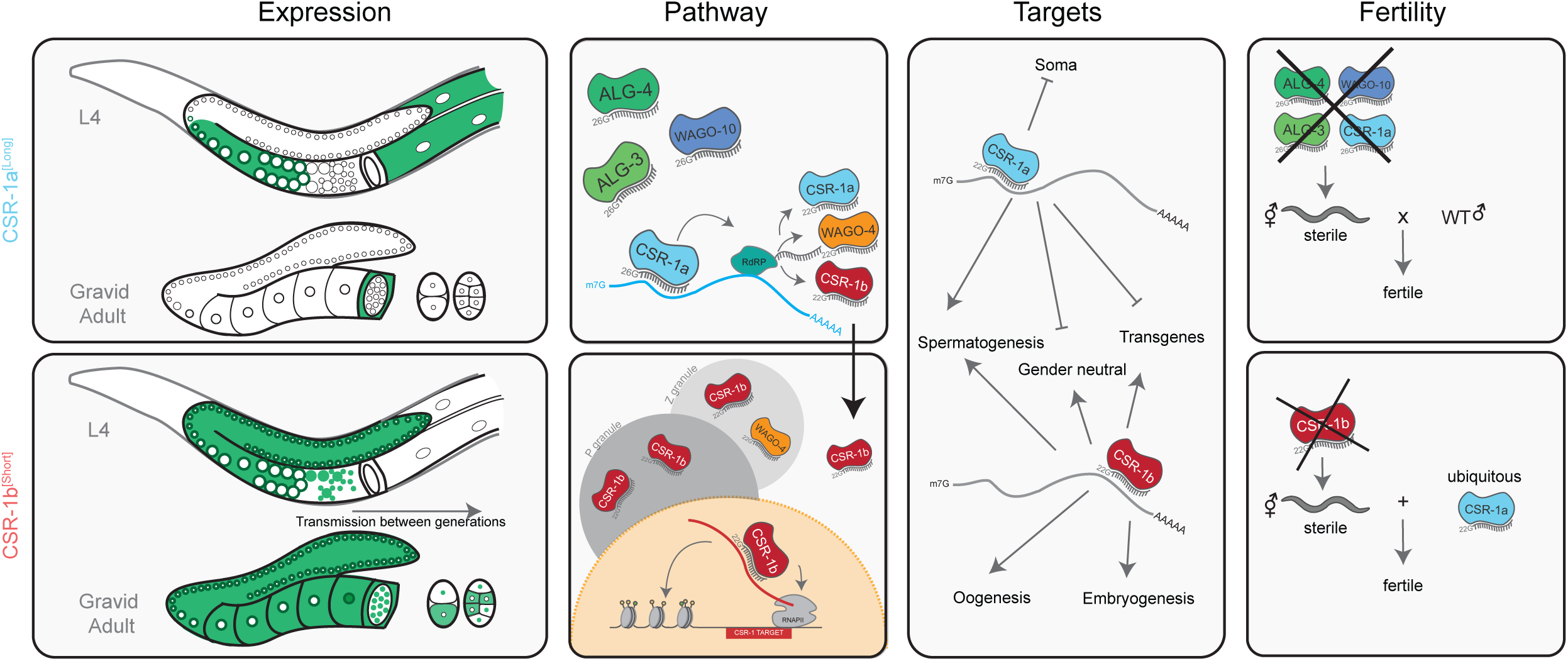
Model for differential functions of CSR-1a and CSR-1b.

### CSR-1 isoforms regulate different facets of fertility

Our previous studies using antibodies that recognize both CSR-1 isoforms failed to distinguish the expression patterns of CSR-1a and CSR-1b (Claycomb et al., 2009). By GFP::3xFLAG tagging the individual CSR-1 isoforms, we found that CSR-1b is expressed constitutively in the germline throughout all stages, and is present in somatic cells of the early embryo, due solely to maternal deposition. In contrast, CSR-1a is expressed in the germline only during spermatogenesis. Moreover, CSR-1a is the only isoform expressed in the adult soma, where it is predominantly present in the intestine and spermatheca. These expression profiles alone shine a light on which CSR-1 isoform may be responsible for which phenotypes. CSR-1b alone carries out essential gene regulatory functions in the hermaphrodite germline and within embryos, where the sterile/lethal phenotype manifests. Therefore, we conclude that *csr-1b* is the essential isoform under “luxurious” lab culture conditions.

Conversely, CSR-1a plays an important role in sperm-based fertility within hermaphrodites. Loss of *csr-1a* alone leads to a transgenerational loss of sperm-based fertility in hermaphrodites. CSR-1a also plays a redundant role with other 26G-RNA associated AGOs during spermatogenesis that is required for male fertility in a single generation at stressful temperatures. Because CSR-1b is also expressed throughout the germline during spermatogenesis, it is conceivable that it too is required for sperm-based fertility. Unfortunately, our data do not specifically address this at this time, and comparing the generation of onset for the transgenerational loss of *csr-1a* fertility to that of *csr-1* null mutants is fraught by the oocyte/embryo fertility/lethality defect. Instead, to assess a role for CSR-1b in sperm-based fertility, we would need to examine male fertility in a transgenerational mating assay similar to (Conine et al., 2013) or rescue the *csr-1(tm892)* null mutant with a *csr-1a* transgene expressed from the *csr-1* promoter. Selective degradation of CSR-1b during spermatogenesis via a degron approach (Martinez et al., 2020) may also enable such an assessment. Related, any male specific roles for either CSR-1 isoform should also be explored in the future. While many aspects of spermatogenesis are consistent between male and hermaphrodite *C. elegans*, there are several features that are sex-specific (L’Hernault, 2009).

### CSR-1 isoform association with germ granules and chromatin

Our previous work showed that CSR-1 co-localizes with PGL-1 and is present within germ granules known as P granules (Claycomb et al., 2009). We initially speculated that CSR-1a might be the only isoform capable of localizing to P granules due to its unique RGG/RG rich N-terminus. RGG/RG motifs have been implicated in RNA binding, recruiting RNA binding proteins, and inducing phase separation *in vitro*—all of which are functions relevant to germ granules (Chong et al., 2018). Loss of *csr-1* has been shown to result in the mis-localization of P granules to the core cytoplasm of the germline, pointing to a role for CSR-1 in the maintenance and proper positioning of P granules (Claycomb et al., 2009; Campbell and Updike, 2015). We observed that both CSR-1 isoforms localize to germ granules, which are likely to be P granules, based on our previous studies (Claycomb et al., 2009). Further, co-localization studies using a strain expressing PGL-1::RFP and GFP::3xFLAG::CSR-1 demonstrated strong overlap in gravid adults, a stage when only CSR-1b is expressed (data not shown).

There are multiple mechanisms by which proteins can associate with phase separated granules, including association with RNA and other proteins that induce phase separation (Chong et al., 2018). The fact that CSR-1b, without an RGG/RG rich domain, and CSR-1a^[G91R]^ localize to germ granules (data not shown) supports the notion that RNA and/or another protein enable the association of CSR-1 with these granules. In addition to its localization to P granules, CSR-1b also appears to associate with the newly identified Z granules (Wan et al., 2018). This would make CSR-1b only the third known component to localize to Z granules (Wan et al., 2018). Perhaps this relationship with Z granules is not surprising, because WAGO-4, which targets a largely overlapping complement of transcripts as CSR-1b via 22G-RNAs, was one of the founding components identified within Z granules (Wan et al., 2018; Xu et al., 2018). Our studies also distinguish germline expressed gene sets that are targeted coordinately by CSR-1b and WAGO-4 (see below). Why these genes are subject to regulation by different AGO/sRNA pathways and how these genes are regulated by those pathways are currently unknown. There is clearly a deeper molecular link between CSR-1b and WAGO-4, P granules and Z granules that demands further study.

We also observed that CSR-1b associates with spermatid and oocyte nuclei and chromatin. These observations are consistent with a previous proteomic study that found CSR-1 peptides enriched on both oocyte and sperm chromatin (Chu et al., 2006), as well as microscopy and ChIP studies that point to an association of CSR-1 with chromatin (Claycomb et al., 2009; Wedeles et al., 2013b; Avgousti et al., 2012; Conine et al., 2013). These collective findings point to a consistent pool of CSR-1b present in the nucleus and/or associated with germ cell chromatin, despite the fact that the bulk of CSR-1b protein is present within the cytoplasm and germ granules (Claycomb et al., 2009). This pool of CSR-1b can be passed from parent to progeny via sperm or oocyte chromatin and presents the possibility for epigenetically marking chromosomes, influencing chromatin landscapes, and reinforcing patterns of transcription in progeny (Gushchanskaia et al., 2019; Conine et al., 2013; She et al., 2009). CSR-1 has been implicated in transgenerational phenotypes, including Mrt and RNAa (Conine et al., 2013; Wedeles et al., 2013b; Seth et al., 2013), and this chromatin association could be but one of several molecular mechanisms by which these phenomena are executed.

### How and why do CSR-1 isoforms associate with different pools of sRNAs?

Based on previous studies of CSR-1 associated sRNAs in males undergoing spermatogenesis and adult hermaphrodites undergoing oogenesis (Conine et al., 2013; Claycomb et al., 2009), we anticipated that CSR-1a and CSR-1b would both associate with 22G-RNAs. Therefore it was remarkable to observe that CSR-1a associates with both 26G-RNAs and 22G-RNAs. Loss of AGOs often leads to destabilization and loss of partner sRNAs (Claycomb et al., 2009; Gu et al., 2009; Conine et al., 2010). A previous study in which total sRNAs were sequenced from *csr-1(tm892)* null mutant males demonstrated that both 22G-RNAs and 26G-RNAs were depleted in this background (Conine et al., 2013). This result was unexpected at the time, but is entirely consistent with our observation that CSR-1a binds to 26G-RNAs.

We were able to rescue the essential functions of *csr-1b* in the oogenic germline and embryo by aberrantly expressing CSR-1a. This result suggests that ectopically expressed CSR-1a is capable of binding CSR-1b 22G-RNAs, as spermatogenesis 26G-RNAs are not produced at this developmental stage (Conine et al., 2010; Vasale et al., 2010), and that ectopic CSR-1a can perform the essential functions of CSR-1b in the oogenic germline. These data support the idea that under certain circumstances, 26G-RNA binding AGOs are also able to associate with 22G-RNAs. Furthermore, when we examined the sRNAs associated with the first-described 26G-RNA AGOs ALG-3, and ALG-4 more closely, we observed a significant proportion of 22G-RNAs enriched in these complexes (ALG-3 targets 555 genes and ALG-4 targets 1082 genes with 22G-RNAs), indicating that this is a feature of multiple 26G-RNA associated AGOs.

How could such plasticity in sRNA binding be achieved? It is likely that accommodating two differently sized sRNAs by one AGO requires conformational changes within the protein. Such conformational changes have been described for other AGOs, such as Drosophila and human miRNA binding AGOs when they are associated with the RISC Loading Complex (Swarts et al., 2014). Therefore, one possibility is that a protein binding partner or complex that associates specifically with CSR-1a assists in discriminating between particular sRNA species. Such a binding partner could be part of a sRNA loading complex, could induce changes in the conformation of the CSR-1a to accommodate longer sRNAs, or could stabilize the association of CSR-1a with 26G-RNAs. In the absence of such a protein co-factor, spermatogenesis AGOs may default to binding the far more abundant 22G-RNAs. We might also expect that this binding partner is co-expressed during spermatogenesis with the AGOs. Post-translational regulation of CSR-1a and ALG-3 and −4 could also play a role in their association with 26G-RNAs or the recruitment of such a cofactor. Although there are many potential post-transcriptional modifications, arginine methylation seems a relevant candidate because CSR-1a, ALG-3 and ALG-4 all possess RGG/RG motifs—which are the sites of methylation (Thandapani et al., 2013) (see below). Finally, localization could be another means to control differences in accessibility to sRNAs or protein co-factors. While CSR-1a, ALG-3 and ALG-4 are enriched in germ granules, they also are found diffusely within the cytoplasm. Additional experiments including mass spectrometry or *in vivo* proximity ligation experiments (Branon et al., 2018; Barucci et al., 2020) will identify the protein binding partners of the 26G-RNA binding AGOs during spermatogenesis. Structural analysis of CSR-1a and ALG-3 and −4 will illuminate how they engage with their sRNA binding partners. Notably, none of the *C. elegans* AGOS have had their crystal structure solved to date. This, coupled with *in vivo* functional analysis of designer *ago* mutants enabled by CRISPR genome editing will shine a light on how these AGOs, and others, carry out their specific molecular roles.

### An extensive network of spermatogenesis sRNA pathways

In addition to CSR1-a, we identified a second novel spermatogenesis-expressed AGO, WAGO-10, and found that CSR-1a, WAGO-10, ALG-3 and ALG-4 all bind to 26G-RNAs that target a core set of 442 spermatogenesis genes. These 442 26G-RNA targets are enriched in phosphatase and phosphate metabolism genes, genes involved in cytoskeleton and pseudopod formation—all critical functions during spermatogenesis and spermiogenesis (L’Hernault, 2006; Raudvere et al., 2019; Ortiz et al., 2014). This overlapping profile of gene targets for the spermatogenesis AGOs pointed to the potential for genetic redundancy. While *csr-1a*, and *alg-3; alg-4* mutants display a transgenerational loss of fertility at high temperatures (Conine et al., 2013), the quadruple spermatogenesis *ago* mutant displayed male-dependent infertility in a single generation at the stressful temperature, supporting that these AGOs function redundantly during spermatogenesis. We did not deeply examine the phenotype of spermatogenesis *ago* mutant sperm, but based on the genes targeted by these pathways, we expect to observe defects in the progression of spermatogenesis and/or in the activation of spermatozoa (Conine et al., 2013; Conine et al., 2010).

By comparing the sets of target genes for CSR-1a, CSR-1b, WAGO-4, WAGO-10, ALG-3, and ALG-4, we were able to stratify spermatogenic germline genes into different gene regulatory modules. While some gene sets are only regulated by WAGO-4-and CSR-1b-associated 22G-RNAs or ALG-3-and ALG-4-associated 26G-RNAs, most are regulated jointly by sRNAs from both classes, with 26G-RNAs presumably acting upstream of 22G-RNAs. A significant proportion of the genes targeted by both 26G-RNAs and 22G-RNAs utilize CSR-1b and WAGO-4. A larger, more heterogeneous group utilizes 22G-RNAs associated with CSR-1b and WAGO-4 or the 22G-RNAs associated with the spermatogenesis AGOs. GO analysis (Raudvere et al., 2019) and manual examination of gene sets involved in various aspects of spermatogenesis and spermiogenesis (White-Cooper and Bausek, 2010; Ellis and Schedl, 2007; L’Hernault, 2006; Chu et al., 2006; Ma et al., 2014) pointed to CSR-1b and WAGO-4 as being important for regulating general germline functions, such as sex determination and maintaining the germline stem cell niche. ALG-3 and ALG-4 along with CSR-1a and WAGO-10 contribute to the regulation of phosphatases that entail a key means by which spermatogenic gene regulation is achieved. These AGOs also regulate genes involved in sperm structure during spermiogenesis and the formation of motile spermatozoa during activation.

Many of the spermatogenesis-enriched genes and sperm proteome genes (Ortiz et al., 2014; Ma et al., 2014) are not well characterized in terms of their homology or function within sperm, which makes it challenging to anticipate the phenotypic implications of sRNA targeting. Moreover, many of these sperm genes are not depleted of sRNAs in *glp-4(bn2)* mutants (Gu et al., 2009) (Fig. S8E-G), despite being the target of sRNAs. This suggests that the sperm genes could also be expressed and targeted by sRNAs in somatic tissues. The link between these tissues and its role in spermatogenesis (Chavez et al., 2018) will be intriguing to explore in the future.

### Somatic gene silencing roles for CSR-1a

CSR-1a is the only CSR-1 isoform expressed in the adult soma, where it appears to silence somatic transgenes and endogenous intestinal transcripts. These activities are most likely mediated through 22G-RNAs, as most 26G-RNAs are germline-specific, and those 26G-RNAs that persist in *glp-4(bn2)* mutants (Gu et al., 2009) do not map to known CSR-1a 26G-RNA target genes (data not shown). In addition to nonsense mutations, we identified missense mutations of specific residues within the RGG/RG repeats (G91R, G120E) of *csr-1a* that led to de-silencing of the repetitive *rpn-2_p_::GFP* transgene in the intestine. Because the rest of the protein is shared between both CSR-1 isoforms, it seems plausible that the only portion of the protein in which we could recover viable mutations is the N-terminus. While the N-terminus of CSR-1a is the least well-conserved portion of the protein, the RGG/RG repeats within this region are nonetheless highly conserved (Fig. S1G).

The specific nature of these missense mutations suggests that the RGG/RG repeats are important for CSR-1a function in the soma, and could point to a role for arginine methylation in regulating the somatic pool of CSR-1a. Arginine methylation is important for the recruitment of tudor domain proteins to AGO, for example, TudorSN and Piwi in Drosophila (Kirino et al., 2009; Liu et al., 2010; Vagin et al., 2009). In the worm, we hypothesize that tudor domain proteins, such as RSD-6 or EKL-1 (a known CSR-1 pathway component) (Claycomb et al., 2009; Gu et al., 2009; Tijsterman et al., 2004; Zhang et al., 2012; Sakaguchi et al., 2014) may interact with CSR-1a in a methyl-arginine dependent manner. Although we did not recover mutations in *rsd-6* from the forward genetic screen, we independently uncovered a role for *rsd-6* along with *csr-1a* alleles in the somatic transgene desilencing assay. Creating the *rsd-6, csr-1a^[G91R]^* double mutant did not enhance the extent of transgene desilencing (data not shown), which might be expected if the G91R mutation disrupted an interaction between RSD-6 and CSR-1a. The G91R mutation also appears to destabilize CSR-1a, which could simply be due to improper folding of CSR-1a^[G91R]^. Alternatively, it is possible that this mutation alters interactions with protein binding partners that are required for CSR-1a stability. At this point, we do not know whether sRNA association with CSR-1a^[G91R]^ is altered, which could also affect CSR-1a somatic functions. We also note that upon recovering the G91R mutation from the screen, we examined its brood size at 25°C and observed a defect (Fig. 2), suggesting that the RGG/RG motifs are important in the germline as well. Nevertheless, the beauty of forward genetics has implicated these residues as important features of CSR-1a, particularly in the soma.

The screen that identified *csr-1a*-specific mutations also recovered alleles of *rde-1*, *rrf-1*, and *rsd-2*. These factors have all been implicated in exogenous and endogenous gene silencing (Sijen et al., 2001; Tabara et al., 1999; Tijsterman et al., 2004). Related to this, *csr-1* was uncovered in early screens for a loss of factors that lead to germline transgene silencing defects, thus the somatic silencing activity of CSR-1a may be linked to a similar role in the germline (Robert et al., 2005). Collectively these factors could entail a novel somatic transgene silencing pathway. Alternatively, we must consider the possibility that these sRNA factors could also endogenously silence the expression of *rpn-2* (via its promoter) and other proteasomal pathway components.

Interestingly, in an attempt to identify other somatic functions for CSR-1a, we also tested whether loss of *csr-1a* resulted in RNAi deficiency (Rde) (Tabara et al., 1999). Null mutants of *csr-1* have been shown to have limited capacity to silence germline transcripts, including *pos-1* and *cdk-1* (Claycomb et al., 2009). However, when we performed RNAi on *dpy-11*, an intestinal-expressed gene, we observed no difference between *csr-1a* mutants and wild-type worms (data not shown). While we were surprised that loss of *csr-1a* did not result in RNAi deficiency, our systematic study of *C. elegans* AGOs points to several other AGOs that are expressed in the intestine and have been previously implicated in RNAi, including the secondary sRNA binding AGOs, SAGO-1 and SAGO-2 are expressed in the intestine (Yigit et al., 2006) (Seroussi, in preparation). One particular interesting candidate is SAGO-2, which has recently been shown to interact with CSR-1a by mass spectrometry (Barucci et al., 2020). Thus, it is possible that these AGOs could function redundantly with CSR-1a to execute exogenous RNAi in this tissue.

Loss of *csr-1a* led to the differential expression of a relatively small fraction of transcripts overall. However, most of these genes displayed increased expression. Moreover, those transcripts that increased showed a stronger correlation with targeting by CSR-1a sRNAs. Because CSR-1a regulates a significant fraction of histone genes, we were concerned about indirect effects on gene regulation. However, the fact that so few genes are differentially expressed in *csr-1a* mutants argues against indirect effects. The mRNA-seq data along with the transgene desilencing assay point to a direct role for CSR-1a in silencing gene expression via its sRNAs, and this role may be important in the soma. Finally, a gene licensing role for CSR-1a has not been ruled out at this point, as there are a number of transcripts that decrease in *csr-1a* mutants, but this dimension of CSR-1a function remains to be explored further.

### Slicer activity of CSR-1 isoforms

Our studies uncover many new dimensions of CSR-1 biology, however they do not address whether Slicer function is important for both CSR-1 isoforms and in what tissues or gene regulatory contexts. Both isoforms possess the catalytic tetrad of residues within the PIWI domain that are required for Slicer activity, and CSR-1 is one of few AGOs in the worm for which Slicer activity has been studied *in vitro* and *in vivo* (Aoki et al., 2007; Gerson-Gurwitz et al., 2016; Fassnacht et al., 2018). Slicer activity has been implicated in fine tuning maternally deposited transcripts with high 22G-RNA density in the embryo (Gerson-Gurwitz et al., 2016). This function can be attributed to CSR-1b based on expression of the isoforms alone. Histone mRNA maturation is another activity for which Slicer activity is anticipated to be required. As both CSR-1 isoforms associate with 22G-RNAs that target half of all histone transcripts, Slicer activity may be important for both isoforms in this context. Clearly, detailed tissue specific studies will be important to delineate the roles of Slicer activity for each isoform throughout development.

### Summary: A tale of two Argonautes

Our studies disentangle a significant amount of complexity surrounding the singly essential *C. elegans* AGO, CSR-1, and generate a myriad of additional questions surrounding the tissue-specific functions of CSR-1 and other AGOs. Newly developed means of expressing and degrading or inactivating proteins in a tissue specific manner will enable us to understand the roles of each CSR-1 isoform at a finer resolution, in specific cells and tissues (Martinez et al., 2020).

In other *Caenorhabditids*, particularly male/female mating species, *csr-1* not only has multiple isoforms, but possesses duplicated gene copies (Tu et al., 2015; Shi et al., 2013). This expansion has been proposed to serve as a defense mechanism against viruses and other invasive nucleic acid transmitted between the members of obligatory mating species (Shi et al., 2013). We have shown that CSR-1a buffers the effects of temperature stress during spermatogenesis to maintain fertility and is involved in gene silencing in the intestine, a tissue that directly interfaces with the environment. Given its roles in *C. elegans*, it is possible that these additional copies of *csr-1* have further specialized to enable other *Caenorhabditis* species to navigate an array of stresses encountered in their natural environment.

Importantly, CSR-1 is not the only Argonaute expressed as multiple isoforms. Several other AGOs, including ALG-1, ALG-2, ERGO-1, and PPW-1, also possess this capacity in *C. elegans*, yet generally only one isoform of each of these proteins has been studied. Our work highlights the importance of studying all AGO isoforms separately, and points to the unexpected new dimensions of tissue-specific sRNA mediated gene regulatory complexity that may be gained by doing so.

## Supporting information

Key Resources Table

## ACKNOWLEDGEMENTS

The authors thank Drs. John Calarco and Arneet Saltzman, and members of the Claycomb lab for critical reading and constructive comments on this manuscript. We thank Dr. Carolyn Phillips and her lab for thoughtful discussions about the CSR-1 isoforms. Thanks to the labs of Drs. Aaron Reinke and Thomas Hurd for use of microscopes, and to WormBase and The Caenorhabditis Genetics Center. We are also grateful to the Toronto Area Worm Community for reagents, protocols, and discussions.

## CONTRIBUTIONS

AGC, NJL, US, MSR, MJA, JW, and AD performed the experiments. AGC, MSR, US, and AD performed computational analyses. JMC, AGC, NJL, and RIM wrote the manuscript. AGC, NJL, MSR, RIM, and AD devised the figures. This work was funded by a Canadian Institutes of Health Research Grant to JMC (PJT-156083). JMC is supported by a CRC Tier II in Small RNA Biology. MSR is supported by a CIHR-CGSM fellowship. Some strains were provided by the CGC, which is funded by NIH Office of Research Infrastructure Programs (P40 OD010440).

## STAR Methods

### RESOURCE AVAILABILITY

#### Lead Contact

Further information and requests for resources and reagents should be directed to and will be fulfilled by the Lead Contact, Julie M. Claycomb

#### Materials Availability

New *C. elegans* strains generated in this study will be deposited in the *Caenorhabditis* Genetics Center, and those that are not deposited are available upon request. All high throughput sequencing data are available through GEO, and all custom data analysis pipelines are available through GitHub (see below). Custom CSR-1 antibodies and any plasmids generated in this study are available upon request.

#### Data and Code Availability

Small RNA-seq and poly(A) selected mRNA-seq data generated in this study are available under the accession number GEO: GSE154678. The *glp-4(bn2)* depleted gene set was obtained from (Gu et al., 2009) available from GEO under accession number GSE18215. Gender neutral, oogenic and spermatogeneic gene lists were defined as in (Ortiz et al., 2014) available from GEO under accession number GSE57109. The WAGO-4 IP/Small RNA sequencing data was obtained from (Xu et al., 2018) available from GEO under accession number GSE112475.

Sperm proteome data was as described (Ma et al., 2014) available at http://159.226.118.206/miaolab/C.elegans%20data.htm. Early Embryo and L4 expression data was obtained from (Gerstein et al., 2010; Spencer et al., 2011) available under accession numbers: SRX004866 and SRX008144.

The custom scripts used to generate the processed data are available at www.github.com/ClaycombLab/Charlesworth_2020.

### EXPERIMENTAL MODEL AND SUBJECT DETAILS

All *C. elegans* strains were derived from the wild type Bristol N2 strain and cultured according to standard growth conditions (Brenner, 1974) on Nematode Growth Medium (NGM) plates inoculated with *E. coli* OP50 at 20°C, unless otherwise stated. All strains used in this study are listed in the Key Resources Table.

### METHOD DETAILS

#### Strain construction and validation

Strains were constructed using the following methods. Genotypes are described in the Key Resources Table.

##### Co-CRISPR

Strains JMC164, GR3042, GR3043 were constructed from JMC101, GE24 [*pha-1(e2123) III]* (Schnabel and Schnabel, 1990) and GR3042 respectively, using the Co-CRISPR technique. All guide RNA constructs were generated by Q5 site directed mutagenesis as described in (Dickinson et al., 2013). Repair template single-stranded DNA oligonucleotides were designed as described in Paix et al., 2014 and Ward, 2015. Injections were performed using editing of *dpy-10* (to generate *cn64* rollers) as a phenotypic co-CRISPR marker (Arribere et al., 2014; Ward, 2015). Injection mixes contained 60ng/μL each of the co-CRISPR and gene of interest targeting the guide RNA/Cas9 construct, and 50ng/μL each of the co-CRISPR and gene of interest repair template oligonucleotides. Guide RNA and homologous repair template sequences are listed in the Key Resources Table and Supplemental Table S7. All CRISPR edits were confirmed by Sanger sequencing.

##### CRISPR/Cas9-triggered homologous with self-excising cassette

CRISPR was performed as described in (Dickinson et al., 2015; Ouyang et al., 2019; Lev et al., 2019). The repair templates were amplified using primers listed in the Supplemental Table S7 and introduced into the vector pDD282 (Addgene #66823) by isothermal assembly (Gibson, 2009). The guide RNA construct was generated with the NEBuilder kit (New England BioLabs, E2621) using the plasmid pDD162 (Addgene #47549) and the primers listed in the Supplemental table S7.

##### miniMos transgenesis

Cloning was performed by isothermal/Gibson assembly (Gibson, 2009). All plasmids used for transgenesis are listed in supplemental table S7. Constructs were assembled in pNL43 (Lehrbach and Ruvkun, 2016). MiniMos injections were performed as described in (Frøkjær-Jensen et al., 2014). In order to make the *csr-1(tm892); csr-1a rescue* strain, exon 1 of *csr-1a* (a 474 bp fragment beginning at the start codon of *csr-1a*) was placed downstream of the *rpl-28* promoter (605 bp immediately upstream of the *rpl-28* start codon) and fused in-frame at the 5′ end of GFP. The 3′ end of the GFP coding sequence was fused in-frame with a DNA fragment containing the remaining exons of *csr-1a* and the *csr-1* 3′UTR (3829 bp beginning at the first codon of exon 2 of *csr-1a*). This construct (assembled in pNL201 and used to generate *mgTi11* and *mgTi12*) drives the ubiquitous expression of CSR-1a^[Long]^ with an internal GFP tag inserted after K158. In order to make the *csr-1(tm892); csr-1b rescue* strain, the DNA fragment containing GFP fused in-frame to *csr-1a* exons 2 to the csr-1 3′UTR (4697 bp, from pNL201) was placed downstream of the *rpl-28* promoter (605 bp immediately upstream of the *rpl-28* start codon). This construct (assembled in pNL213 and used to generate *mgTi13)* drives the ubiquitous expression of CSR-1b^[Short]^ fused to GFP at its N-terminus via a short linker with the sequence ‘GLNSD’.

##### Generation of the rpn-2_p_::gfp transgene

The integrated *mgIs75*[*rpn-2_p_::gfp::unc-54*] transgene was generated from *sEx10255* (Hunt-Newbury et al., 2007). EMS mutagenesis was used to induce integration of the extrachromosomal array.

#### Developmental staging

Synchronised worms were reared on fresh seeded plates at 20°C and harvested as follows: L1s at 8 hours after plating, L2s at 26 hours, L3s at 34.5 hours, L4s at 48 hours, young adults at 60 hours and gravid adults at 72 hours. Embryos were collected from gravid adult worms 72 hours after plating.

#### Mutagenesis screening and genome sequencing

Mutagenesis was performed by treatment of L4 animals in 47mM EMS for 4 hours at 20°C. Genomic DNA was prepared using the Gentra Puregene Tissue kit (Qiagen, #158689) according to the manufacturer’s instructions. Genomic DNA libraries were prepared using the NEBNext genomic DNA library construction kit (New England Biololabs, #E6040), and sequenced on an Illumina Hiseq instrument. Deep sequencing reads were analyzed using Cloudmap (Minevich et al., 2012).

#### Immunofluorescence microscopy

Gonads and embryos were excised in 1xPBS with 0.5mM Levamisole on poly-L-lysine coated slides from adult or gravid hermaphrodites, respectively, grown at 20°C, then frozen on dry-ice for at least 10 minutes or stored at −80°C for up to two weeks. Fixation and incubation with primary and secondary antibodies were performed as described by (Claycomb et al., 2009). DNA was stained with 1μg/ml DAPI in 1xPBS. Imaging was performed on a Nikon TiE inverted microscope and Nikon C2 confocal system with a PL Apochromat Lambda 60x Oil Immersion objective. Images were acquired using NIS Elements AR software and processed using ImageJ software (Schneider et al., 2012).

#### Live fluorescent microscopy

Staged animals were picked in a 1x M9 droplet with 2μL 2mM Levamisole on standard microscopy slides with 2% agarose pads and covered with 22mm coverslips. Imaging was performed on a Nikon TiE inverted microscope and Nikon C2 confocal system with a PL Apochromat Lambda 60x Oil Immersion objective. Images were acquired using NIS Elements AR software. For time course images, larval stage worm and embryo images were taken every 60 seconds. All images were acquired with the same settings and exported via NIS-Elements AR Software. Any post-acquisition adjustments made to images were applied equally between control and experimental sets using Fiji (Schindelin et al., 2012) and Nikon NIS viewer.

#### Imaging released sperm

Males were moved to male-only plates 1-2 days prior to the assay in order to help maximize sperm retention. 2mL of 1X Sperm buffer (50mM Hepes, 25mM KCl, 35mM NaCl, 1mM MgSO_4_, 5mM CaCl_2_, 10mM Dextrose/D-Glucose, pH 7.8, filter sterilized) was added to a crystallization dish with 0.5μL of 5μg/ml DAPI. After letting the males soak for 5 minutes in the buffer, they were transferred to poly-L-lysine coated slides and any excess buffer was removed by wicking with filter paper. Once worms were adequately anesthetized with 2mM levamisole, they were dissected to release sperm/germline by making a small cut at the tail end with a 17-gauge needle. A 22mm coverslip was placed on top and samples were imaged immediately using Sequential Scan Mode on a Leica SP8 confocal microscope with a 63x oil immersion objective.

#### CSR-1 zygotic expression assay

Male worms homozygous for GFP::3XFLAG::CSR-1 were crossed to N2 hermaphrodites. After 24 hours hermaphrodites were collected and dissected to acquire early embryos. To acquire later staged embryos and young larvae, plates were washed off with 1X M9 into microcentrifuge tubes and transferred to slides with 2% agarose pads. 2μL of Levamisole was added to immobilize larvae. Samples were imaged at 60x on a Nikon TiE inverted microscope and Nikon C2 confocal system utilizing the NIS-Elements AR Software.

#### Imaging analysis of rpn-2_p_::gfp transgenic worms

Bright field and GFP fluorescence images showing *rpn-2_p_::gfp* expression were collected using a Zeiss AxioZoom V16, equipped with a Hammamatsu Orca flash 4.0 digital camera, using Axiovision software, and were processed using ImageJ software (Schneider et al., 2012). Images shown within the same figure panel were collected together using the same exposure time and then processed identically in ImageJ. To quantify *rpn-2_p_::gfp* expression, the maximum pixel intensity within the intestine of each animal was measured using ImageJ. Values were then normalized such that wild type expression is equal to 1.

#### Brood Size Counts

All brood size assays were conducted on N2, tagged strains and mutants reared at 20°C and 25°C. Parents were propagated at 20°C and 10-20 L4 individuals (1 worm/plate) were shifted to the temperature at which the assay was performed. Every 24 hours the adult was transferred to a fresh plate and embryos were counted. The number of larvae was scored the following day. This continued until the adult hermaphrodite was no longer laying eggs. The total number of embryos laid by a single hermaphrodite over the course of the egg-laying period was determined (total brood size), as well as the total number of larvae that hatched (viable brood size). Plots and statistical analysis were performed with GraphPad Prism.

#### Male mating rescue assay

N2 and mutant hermaphrodites were reared at 20°C. Ten L4 hermaphrodites of each assayed strain were singled out onto plates with either five N2 males or alone. Worms were maintained at 25°C for 48 hours and embryos/oocytes were imaged on a Leica S9 i Stereomicroscope with a 10MP CMOS camera. At 60 hours post temperature shift plates were assessed for the presence of new males to indicate successful mating and successful plates were imaged again in the same manner.

#### Protein lysate preparation

Synchronous populations of animals were grown at a density of 90,000-100,000 animals per 15cm Petri dish on NGM with OP50 *E.coli* as food source and harvested at staged time points in 1x M9 buffer (22mM Potassium dihydrogen phosphate, 42mM Disodium phosphate, 86mM Sodium chloride) and washed with ddH_2_O. The worm pellets were flash frozen in a dry ice/ethanol bath and stored at −80°C. The frozen pellets were resuspended 1:1 (v/v) in ice-cold DROSO complete buffer (30mM Hepes, 100mM Potassium Acetate, 2mM Magnesium Acetate, 0.1% Igepal CA 630 (Sigma-Aldrich), 2mM DTT, 4x concentration Complete Protease Inhibitor (Roche) and 1% (v/v) Phosphatase Inhibitors 2 and 3 (Sigma-Aldrich) or EDTA Complete buffer (30mM Hepes, 100mM Potassium Acetate, 10mM EDTA, 10% Glycerol, 0.1% NP-40 or Igepal CA630 (Sigma-Aldrich), 2mM DTT, 1 tablet/5mL Protease inhibitor (Roche), 1:100 Phosphatase inhibitor 2 (Sigma-Aldrich), 1:100 Phosphatase inhibitor 3 (Sigma-Aldrich) and homogenized using stainless steel dounces (Wheaton Incorporated) until worms and embryos were no longer visible under the microscope. In cases where RNA was to be isolated from immunoprecipitations using the protein lysates, 1% (v/v) SUPERaseIn RNase Inhibitor (ThermoFisher) was added to the buffer prior to homogenization. Lysates were centrifuged at 13,000xg for 10min at 4°C, and the supernatant transferred to a fresh tube. The BioRad Lowry assay kit was used to determine total protein concentration using a Nanodrop 1800C spectrophotometer prior to use in subsequent assays.

#### Western blotting

Samples were resolved by SDS-PAGE on precast gradient gels (4-12% Bis-Tris Bolt gels, ThermoFisher) and transferred to Hybond-C membrane (Amersham Biosciences) using a Bio-Rad semi-dry transfer apparatus (25V for 45min). After washing with PBST (0.1% (v/v) Tween-20 in PBS) and blocking with 5% milk-PBST (137mM Sodium Chloride, 10 mM Phosphate, 2.7mM Potassium Chloride, pH 7.4, and 5% w/v dried nonfat milk), the membrane was then incubated overnight at 4°C in primary antibodies (Key Resources Table). After 3x 10min washes in PBST, the membrane was blocked once again with 5% milk-PBS-T for 1 hour at room temperature, then incubated for 1 hour with secondary antibodies conjugated to HRP (Cell Signaling Technology), washed 3 x 5min in PBST and visualized using Luminata Forte Western HRP substrate (Sigma-Aldrich). Primary antibodies for FLAG [1mg/ml], CSR-1 (antibody 49C1) [0.25 mg/ml] were used at 1:3000 and CSR-1a (antibody 55) [0.57 mg/ml] was used at 1:500. Secondary antibodies listed in the Key Resources table [1mg/ml] were used at a concentration of 1:1000.

#### Immunoprecipitation

For each IP and co-IP experiment, 5mg total protein lysate was used per reaction. For small RNA cloning 3 to 5 x5mg IPs were performed and beads were pooled at the final wash step. GFP-Trap Magnetic Agarose beads (ChromoTek) equilibrated in EDTA Complete buffer were used for anti-GFP IPs and RFP-Trap beads were used for the Mock IP control. Dynabead Protein G beads (Invitrogen) were used for all non GFP/RFP IPs in the manuscript. CSR-1 rabbit peptide antibodies used for IPs were 49C2 [1.25 mg/ml] for total CSR-1 and 55 [0.57 mg/ml] for CSR-1a (Claycomb et al., 2009). Dynabeads were resuspended in PBS with 0.1% Tween and incubated while rotating with 5μg of CSR-1 antibody (or rabbit IgG for the Mock IP control) for 10min at RT, before being washed with EDTA complete buffer. Samples were incubated with 25μL of bead slurry for 1 hour while rotating at 4°C. Samples were then washed with EDTA Complete buffer 5x 10min each and when multiple IPs were done then beads were combined in one sample tube. Protein was eluted from the beads and denatured by incubation with 4x sample buffer (4X Bolt Buffer - ThermoFisher) and 10X reducing agent for 10min at 70°C. Input samples were prepared from the same lysates at a concentration of 2mg/ml using ThermoFisher 4X Bolt Buffer and 10x reducing agent. For samples being used for library construction, 10% of the sample was taken for western blot to determine efficacy of the IP. The remaining sample was mixed with 4x Tri-Reagent (Molecular Research Center, Inc. or Sigma-Aldrich) and extracted as described below.

#### RNA sample preparation

Synchronized staged animals were collected in 1X M9 and washed in M9 and MilliQ water. Upon adding 4x volume Tri-Reagent (Molecular Research Center, Inc. or Sigma-Aldrich), samples were flash frozen in a dry ice/ethanol bath and stored at −80°C for later RNA extraction. Total RNA extraction and ethanol precipitation were performed as described in (Claycomb et al., 2009), followed by quantification using a Nanodrop 1800C spectrophotometer.

#### Small RNA library construction

RNA for sRNA libraries, was isolated from either total worms or IP/ Input samples as described above without DNAse treatment. Maximum 4μg of RNA from the IP samples were treated with 5′ Polyphosphatase (Lucigen) for 30min at 30°C and cleaned following a standard phenol:chloroform:isoamyl alcohol and ethanol precipitation. 1μg of total RNA from each sample was used for sRNA library preparation using the NEBNext Small RNA Library Prep Set for Illumina (New England Biolabs). The resulting PCR product DNA of ∼150bp was visualized using 10% PAGE stained with SYBRGold (ThermoFisher) and the bands of interest were excised. DNA was eluted overnight at room temperature in a buffer containing 3.3M NaCl, 100mM Tris and 1mM EDTA, and precipitated with 20μg glycogen as the carrier and 1X volume isopropanol overnight at −20°C. The resulting DNA was resuspended in 20μL ultrapure water. And quantified using a Qubit HS RNA kit (ThermoFisher). All samples were pooled in equal amounts into a 20nM solution and sequenced at 50bp single end reads on an Illumina HiSeq 2500 Sequencing System. For each sample, two biological replicates were prepared.

#### mRNA library construction

RNA was isolated from whole worms as described in RNA sample preparation above. 2μg of each sample was treated with DNase I (Ambion), phenol:chloroform extracted, ethanol precipitated, resuspended in Ultrapure™ Distilled Water and quantified using a Qubit HS RNA kit (ThermoFisher). 1μg of total DNase treated RNA was used for library preparation using the NEBNext Ultra II Directional RNA Prep Kit for Illumina - with Poly-A selection, according to the manufacturer manual and samples were quantified the Qubit HSDNA kit (ThermoFisher). All samples were pooled in equal amounts into a 20nM solution and sequenced 50bp single end reads on a Illumina HiSeq 2500 Sequencing System. For each genotype and temperature, three biological replicates were prepared.

### DATA ANALYSIS

#### Small RNA Enrichment Analysis

Raw read quality was assessed via FASTQC and adapters were trimmed via Cutadapt (Martin, 2011) using the adapter sequence “AGATCGGAAGAGCACACGTCTGAACTCCAGTCAC”. Any reads which were not trimmed and were outside of the 16-30 nucleotide range were discarded. (Except for the CSR-1 Antibody IP and corresponding Input sample, reads within the 16-27 nucleotide range were used.) Reads were then mapped to the *C. elegans* genome (WormBase release WS262) using STAR-aligner (Dobin et al., 2013). A custom script was used to count the number of reads for each feature (e.g. protein coding genes, pseudogenes, etc.). Enrichment of sRNA for each feature was determined using a custom analysis script. Enriched genes have a minimum of 5 RPM in both IP samples, and >2 log_2_ fold enrichment in both IPs relative to Input samples. A custom script was used to generate the size and first nucleotide distribution plots. The distribution and enrichment scatter plots were made using the ggplot2 package (Wickham, 2016). Venn plots were made using VennDiagram package (Chen and Boutros, 2018). The overlap between enriched sRNAs and various published gene sets was determined and plotted using the VennDetail Package (Guo and McGregor, 2020) using percentage parameters. Additional genes and gene groups of interest were examined using WormMine (Smith et al., 2012; Kalderimis et al., 2014), WormExp (Yang et al., 2016) and g:Profiler (Raudvere et al., 2019). The significance in overlap between gene sets was calculated using Fisher’s Exact Test. Untemplated nucleotide additions to sRNA 3′ ends were identified and quantified using a custom script. Metagene plots of sRNA distributions along target genes were made with the Metagene2 package (Fournier et al., 2020). Replicates were aggregated and normalized by RPM. Standard package parameters were used. For clustering diagrams, sRNA targets were grouped via hierarchical clustering by euclidean distance using the pheatmap package (Kolde, 2019). All custom analysis scripts mentioned above are available at www.github.com/ClaycombLab/Charlesworth_2020.

#### mRNA Differential Gene Expression Analysis

mRNA data quality was initially assessed via FASTQC, and any adapter sequences were trimmed via Cutadapt (Martin, 2011) using the adapter sequence “AGATCGGAAGAGCACACGTCTGAACTCCAGTCAC.” All reads without adapters were retained for analysis. Reads were then mapped to the *C. elegans* genome (WormBase release WS262) using STAR-aligner (Dobin et al., 2013). The number of reads mapping to each gene feature was determined using summarizeOverlaps from the GenomicAlignment package (Lawrence et al., 2013). Pre-filtering was done to remove all rows possessing only a single count or less across all samples. Differential expression and statistical analysis were performed using DESeq2 (Love et al., 2014) and significantly-changed genes were designated by the threshold of >2-fold change (+/-) and <0.01 *p_adj_*. Data were then plotted using the EnhancedVolcano package (Blighe and Rana, 2020). Any genes that were also differentially expressed in wild-type (N2) samples due to temperature were manually removed. Overlap between differentially expressed mRNAs and published gene sets was determined and plotted using the VennDetail Package (Guo and McGregor, 2020) using percentage parameters. The significance in overlap between gene sets was calculated using Fisher’s Exact test. Additional genes and gene groups of interest were examined using WormMine (Smith et al., 2012; Kalderimis et al., 2014), WormExp (Yang et al., 2016) and g:Profiler (Raudvere et al., 2019).

#### Gene Ontology Analysis

Gene lists generated via our custom pipelines were subjected to Gene Ontology Analysis using g:Profiler (Raudvere et al., 2019) under default settings.

#### Sequence Alignment and Cladogram Construction

*C. elegans csr-1* homologous sequences for other *Caenorhabditis* species were obtained from the NCBI database (Benson et al., 2017), then aligned with MUSCLE (Edgar, 2004) and custom coloured to highlight particular amino acids relevant to this study in SeaView (Gouy et al., 2010). The cladogram was constructed based on the alignment.

#### Quantification and Statistical Analysis

Statistical analyses of brood sizes were conducted using GraphPad Prism (v. 8.4.1). Comparison of only two samples was calculated using an unpaired *t*-test with an alpha of 0.05. Comparisons of more than two samples were calculated using ordinary one-way ANOVA, followed by Tukey’s multiple comparisons test.

Statistical analysis for all Venn-pie Gene set enrichment comparisons was calculated using Fisher’s Exact Test.

sRNA-seq data were determined to be enriched by the presence of >5RPM in each IP replicate, and >2 log_2_ fold increase in the number of reads versus Input. Similarly, sRNA-seq data were determined to be depleted (Gu et al., 2009) by the presence of >5RPM in Input, and >2 log_2_ fold decrease in the number of reads in the mutant versus Input.

All sRNA samples, including those published by other groups were analyzed using our own computational pipeline for consistency.

mRNA-seq data samples were determined to be differentially expressed by the presence of >1 read per feature, >2-fold (+/-) change in expression and *p_adj_* of <0.01.

---

**Supplemental Figure 1.**
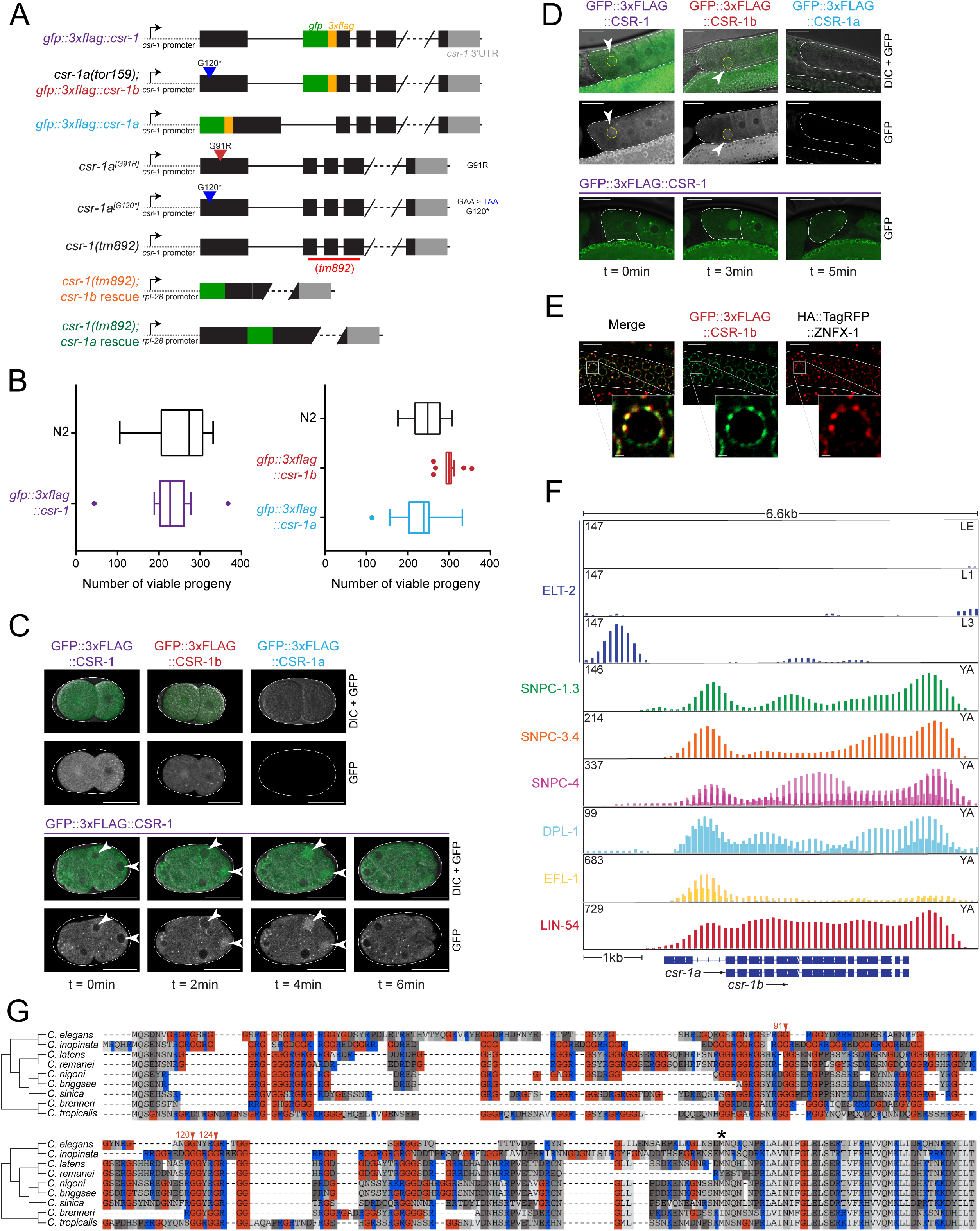
Additional characterization of GFP::3xFLAG::CSR-1a and GFP::3xFLAG::CSR-1b [Related to Fig. 1]. **A)** Schematic representation of all *csr-1* mutant strains, endogenously tagged strains and single copy transgene insertion strains used in this study. **B)** Box plot representation of viable brood size counts for the strains expressing endogenously tagged CSR-1 compared to wild type, N2. Median is represented, with Tukey whiskers. Significance was determined with a one-way ANOVA (alpha=0.05) with Tukey’s multiple comparison test. Differences are not significant unless marked. N for each genotype is 14 or more individual P_0_ hermaphrodites. **C)** Fluorescence micrographs showing nuclear localization of endogenously tagged CSR-1 isoforms in 2-cell embryos during nuclear envelope breakdown, and time lapse of nuclear localization of endogenously tagged CSR-1 in a developing 8-cell embryo over the course of six minutes. Scale bar, 25μm. **D)** Fluorescence micrographs showing localization of endogenously tagged CSR-1 isoforms in oocytes and time lapse of endogenously tagged CSR-1 localization in the −1 oocyte over the course of five minutes. Scale bar, 25μm. **E)** Fluorescence micrographs showing GFP::3xFLAG::CSR-1b colocalization with HA::TagRFP::ZNFX-1 (a Z granule marker (Wan et al., 2018)) to germ granules. Scale bar, 10μm. Inset scale bar, 1μm. **F)** Notable transcription factors enriched in binding within the promoters of each *csr-1* isoform, from the ModERN project (Kudron et al., 2018). **G)** Multiple sequence alignment of the N-terminal region of *csr-1* for nine *Caenorhabditis* species. Arrows indicate *C. elegans* amino acid positions mutated as a result of the forward genetic screen (as shown in Figure 6). The asterisk indicates the first amino acid of CSR-1b in *C. elegans*.

**Supplemental Figure 2.**
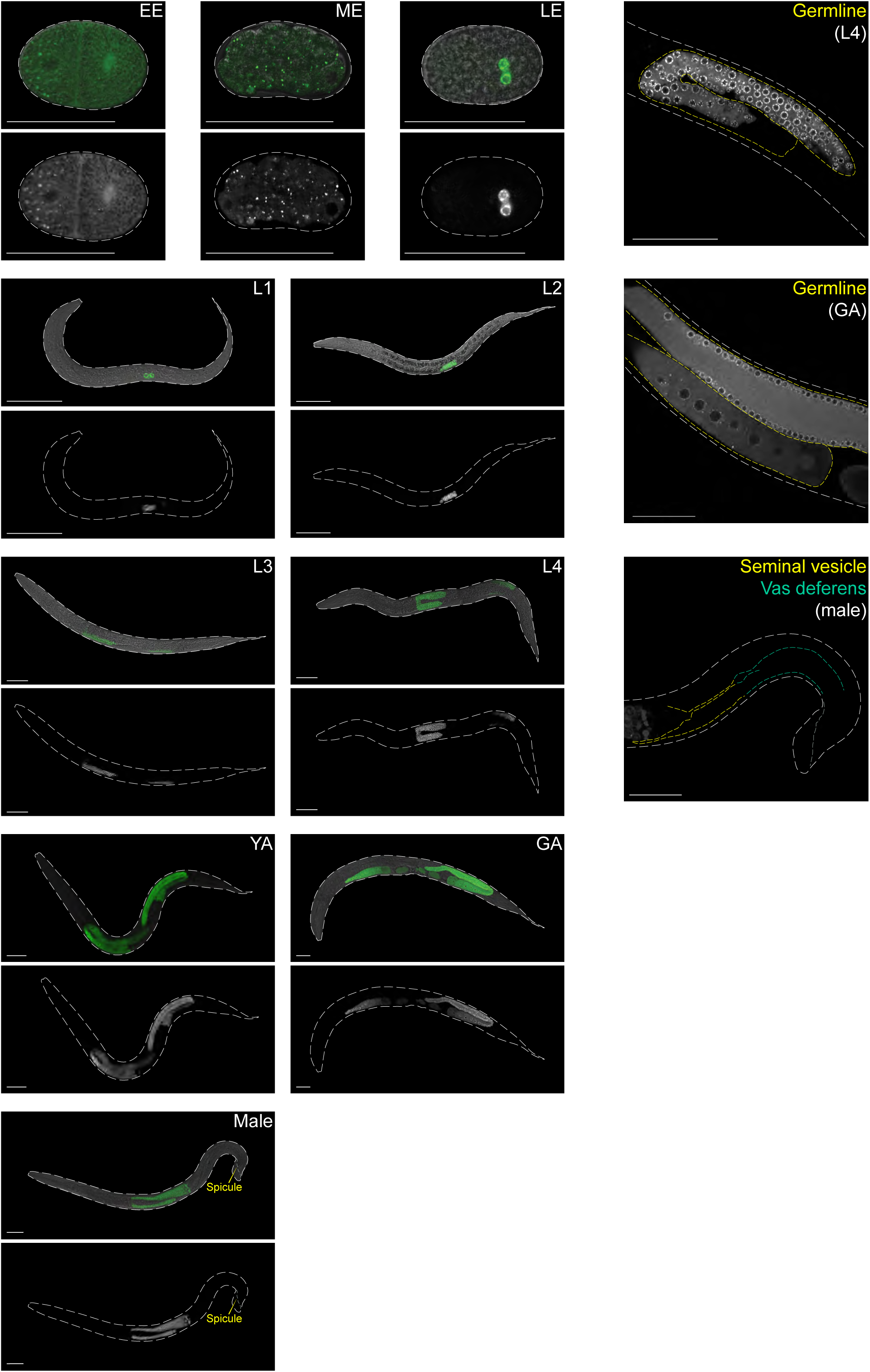
Fluorescence micrographs of whole animals expressing GFP::3xFLAG::CSR-1b throughout development [Related to Fig. 1]. (Hermaphrodite stages: EE=early embryo, ME=mid embryo, LE=late embryo, L1-L4=larval stage 1-4, YA=young adult, GA=gravid adult) and adult male, with close-up images of L4 germline, GA germline and male somatic gonad, respectively. Scale bars, 50μm.

**Supplemental Figure 3.**
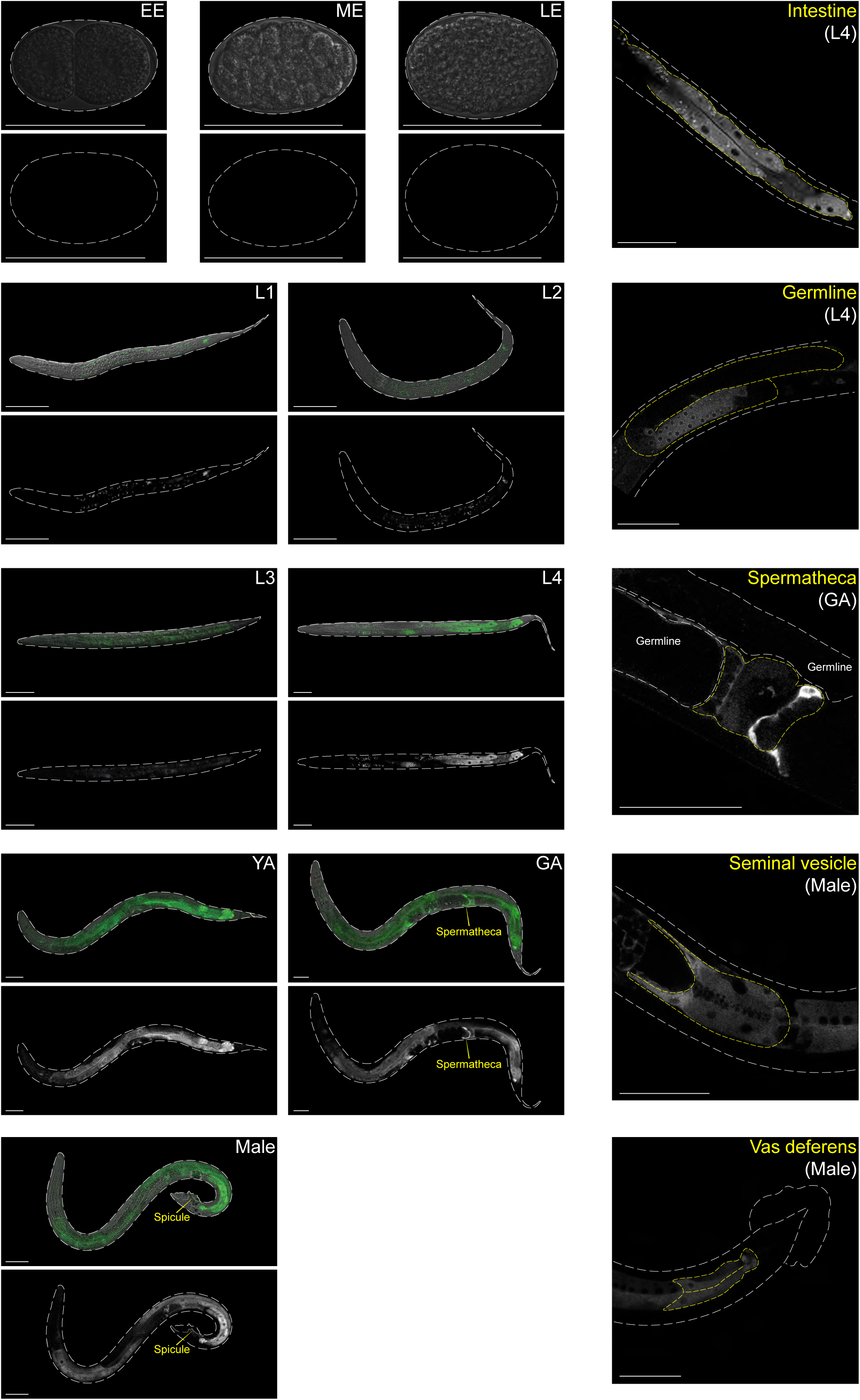
Fluorescence micrographs of whole animals expressing GFP::3xFLAG::CSR-1a throughout development [Related to Fig. 1]. (Hermaphrodite stages: EE=Early Embryo, ME=Mid Embryo, LE=Late Embryo, L1-L4=Larval stage 1-4, YA=Young Adult, GA=Gravid Adult) and adult male, with close-up images of L4 intestine, L4 germline, GA spermatheca, male seminal vesicle and male vas deferens, respectively. Scale bars, 50μm.

**Supplemental Figure 4.**
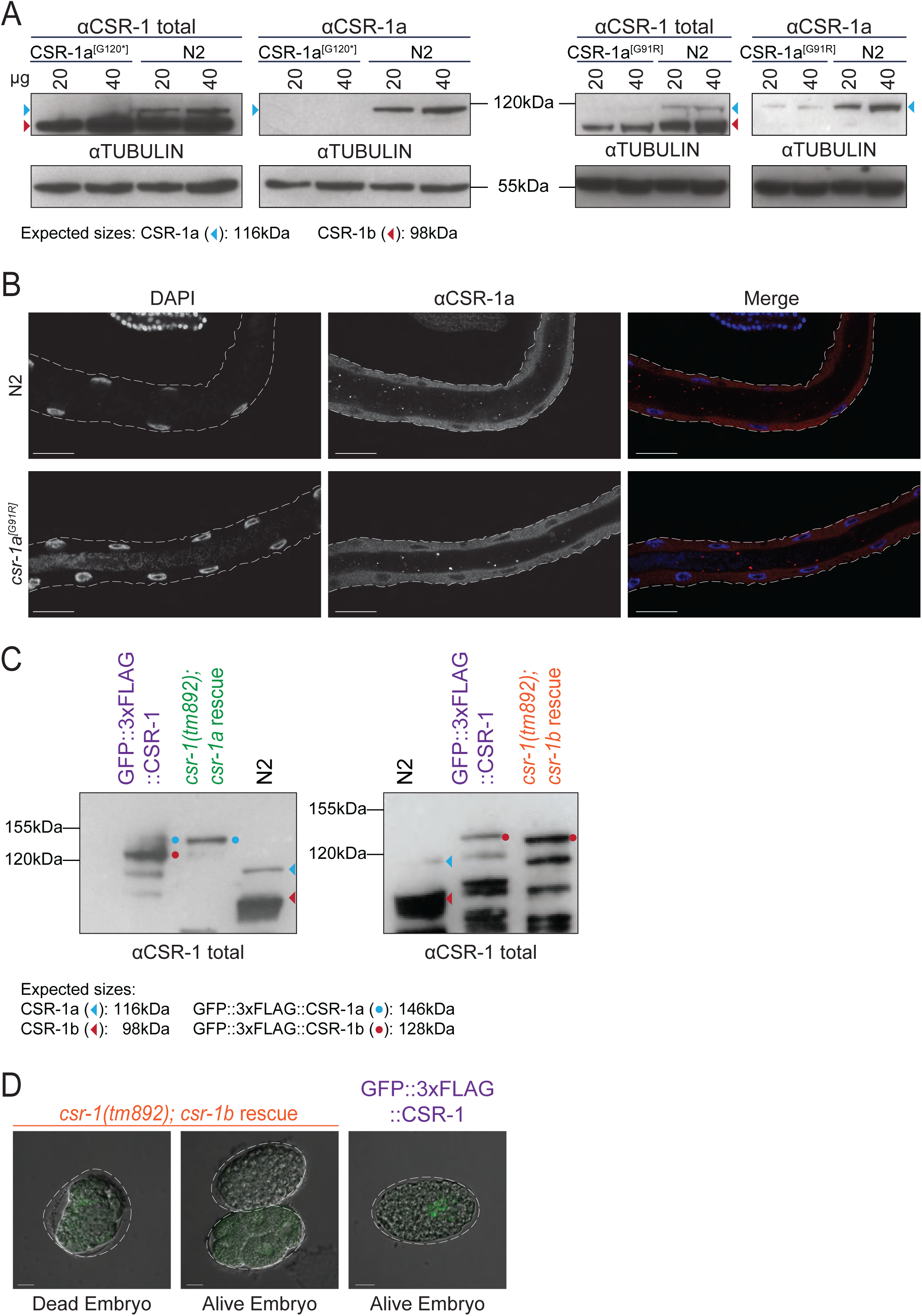
Additional characterization of *csr-1* mutants and rescued strains [Related to Fig. 2]. **A)** Western blots of CSR-1a^[G120*]^, CSR-1^[G91R]^ and N2 lysate probed with antibodies against both isoforms (Antibody 49C2) and CSR-1a only (Antibody 55) with alpha-tubulin as a loading control. **B)** Fluorescence micrographs of dissected intestines of N2 and CSR-1a^[G91R]^ stained with DAPI and an antibody against CSR-1a (Antibody 55). Scale bar, 25μm. **C)** Western blot of endogenously tagged rescued CSR-1 strains and N2 probed with antibody against both CSR-1 isoforms (Antibody 49C2). **D)** Fluorescence micrographs of embryos of the *csr-1(tm892); csr-1b* rescue depicting both dead and alive embryos, compared to embryos of GFP::3xFLAG::CSR-1 animals. Scale bar, 10μm.

**Supplemental Figure 5.**
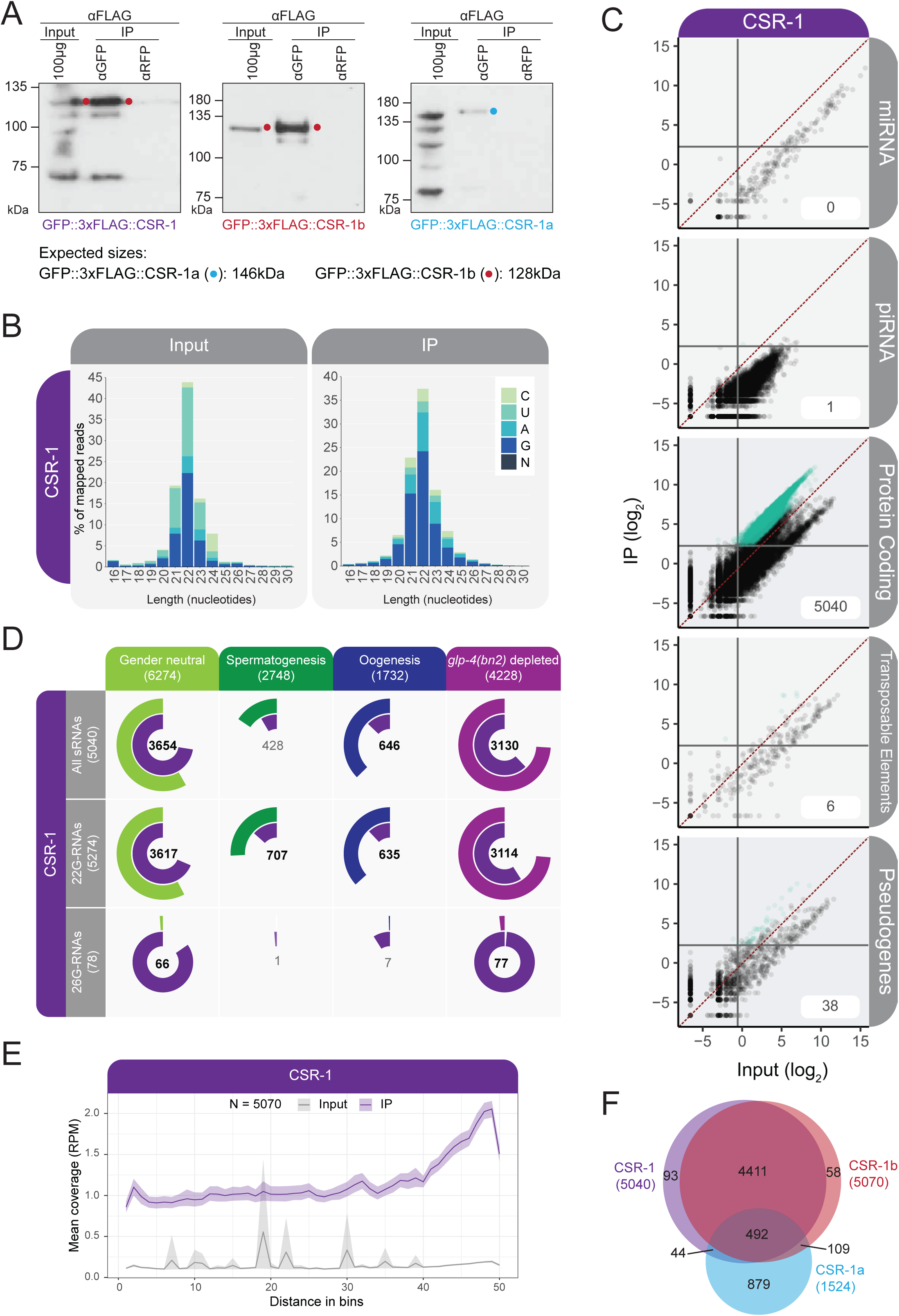
Small RNA analysis of GFP::3xFLAG::CSR-1 [Related to Fig. 3]. **A)** Western Blot using FLAG antibodies for detecting GFP::3xFLAG::CSR-1, GFP::3xFLAG::CSR-1b and GFP::3xFLAG::CSR-1a, respectively, from IPs used for sRNA library preparation. **B)** Bar graph of first nucleotide and size distribution of normalized reads from Inputs and IPs of GFP::3xFLAG::CSR-1. **C)** Scatter plot of sRNA data showing enrichment in IP over Input. Teal dots represent those genes/transcripts which have a minimum of 5 RPM in IP samples and are 2-fold enriched relative to Input in both replicates. **D)** Venn-pie diagrams show comparisons between IP-enriched sRNAs which are antisense to protein coding genes (inner circle) and germline gene sets from Fig. 3 (Ortiz et al., 2014; Gu et al., 2009) (outer circle) All sRNAs = no size restriction of sRNAs, 22G-RNAs = sRNAs in the range of 21-23nt and 26G-RNAs = sRNAs in the range of 25-27nt. The size of each color-coded circle represents the proportion of each sample that is shared between the two sets. Numbers in the center of the circle indicate the number of genes that overlap between the two data sets. Numbers in bold indicate statistically significant overlap as calculated by Fisher’s Exact Test, *p*<0.001. **E)** Metagene plot of the distribution of enriched sRNAs antisense to protein coding genes along the gene body. Number of genes represented is denoted at the top as N. **F)** Venn diagram depicting the overlap in protein coding gene targets of the All sRNAs class enriched in each of the CSR-1 IPs.

**Supplemental Figure 6.**
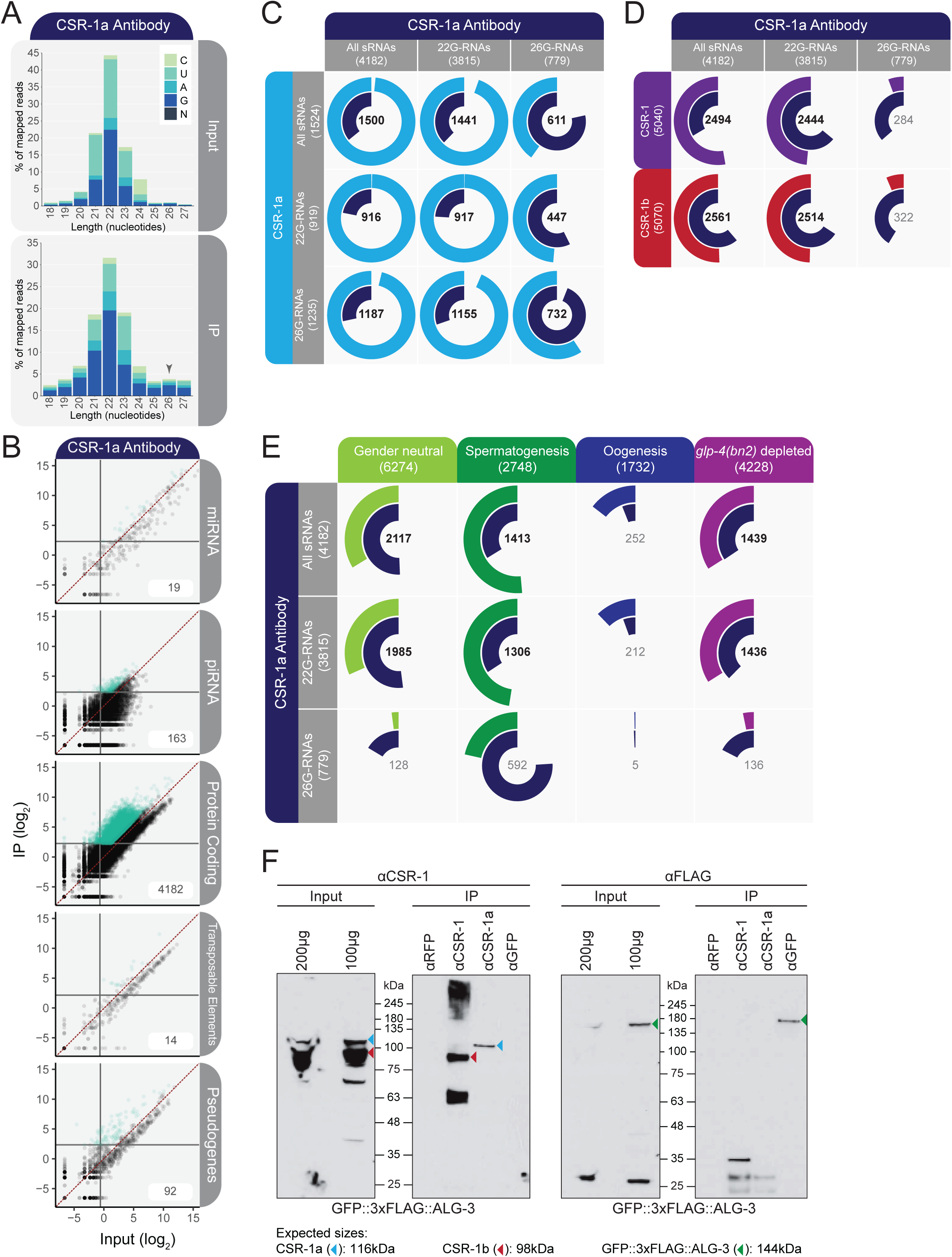
Small RNA analysis of CSR-1a antibody IP [Related to Fig. 3]. **A)** Bar graph of first nucleotide and size distribution of normalized reads from Inputs and IPs of N2 worms with a CSR-1a-specific peptide antibody. Arrow highlights 26G-RNAs. **B)** Scatter plot of sRNA data showing enrichment in IP over Input. Teal points represent those which have a minimum of 5 RPM in IP samples and are 2-fold enriched relative to Input in both replicates. **C-D)** Venn-pie diagrams showing the comparison between IP samples for enriched sRNAs antisense to protein coding genes (GFP IP: inner circle; CSR-1a antibody: outer circle). For CSR-1a: All sRNAs = no size restriction of sRNAs, with no first nucleotide bias, 22G-RNAs = sRNAs in the range of 21-23nt, with no first nucleotide bias, 26G-RNAs = sRNAs in the range of 25-27nt, with no first nucleotide bias. For CSR-1 and CSR-1b, All sRNAs are used. The size of each color-coded circle represents the proportion of each sample that is shared between the two sets. Numbers in the center of the circle indicate the number of genes that overlap between the two data sets. Numbers in bold indicate statistically significant overlap as calculated by Fisher’s Exact Test, *p*<0.001. **E)** Venn-pie diagrams showing comparisons between CSR-1a Antibody IP-enriched sRNAs antisense to protein coding genes (inner circle) and germline gene sets from Fig. 3 (Ortiz et al., 2014; Gu et al., 2009) (outer circle). The size of each color-coded circle represents the proportion of each sample that is shared between the two sets. Numbers in the center of the circle indicate the number of genes that overlap between the two data sets. Numbers in bold indicate statistically significant overlap as calculated by Fisher’s Exact Test, *p*<0.001. **F)** IP-Western blots of GFP::3xFLAG::ALG-3 with endogenous CSR-1. IPs are performed as designated (anti-CSR-1 recognizes both isoforms, anti-CSR-1a recognizes only CSR-1a, and anti-GFP recognizes GFP::3xFLAG::ALG-3). Blots were probed with anti-CSR-1 and anti-FLAG. Expected sizes: CSR-1a: 116kDa, CSR-1b: 98kDa, GFP::3xFLAG:ALG-3: 144kDa.

**Supplemental Figure 7.**
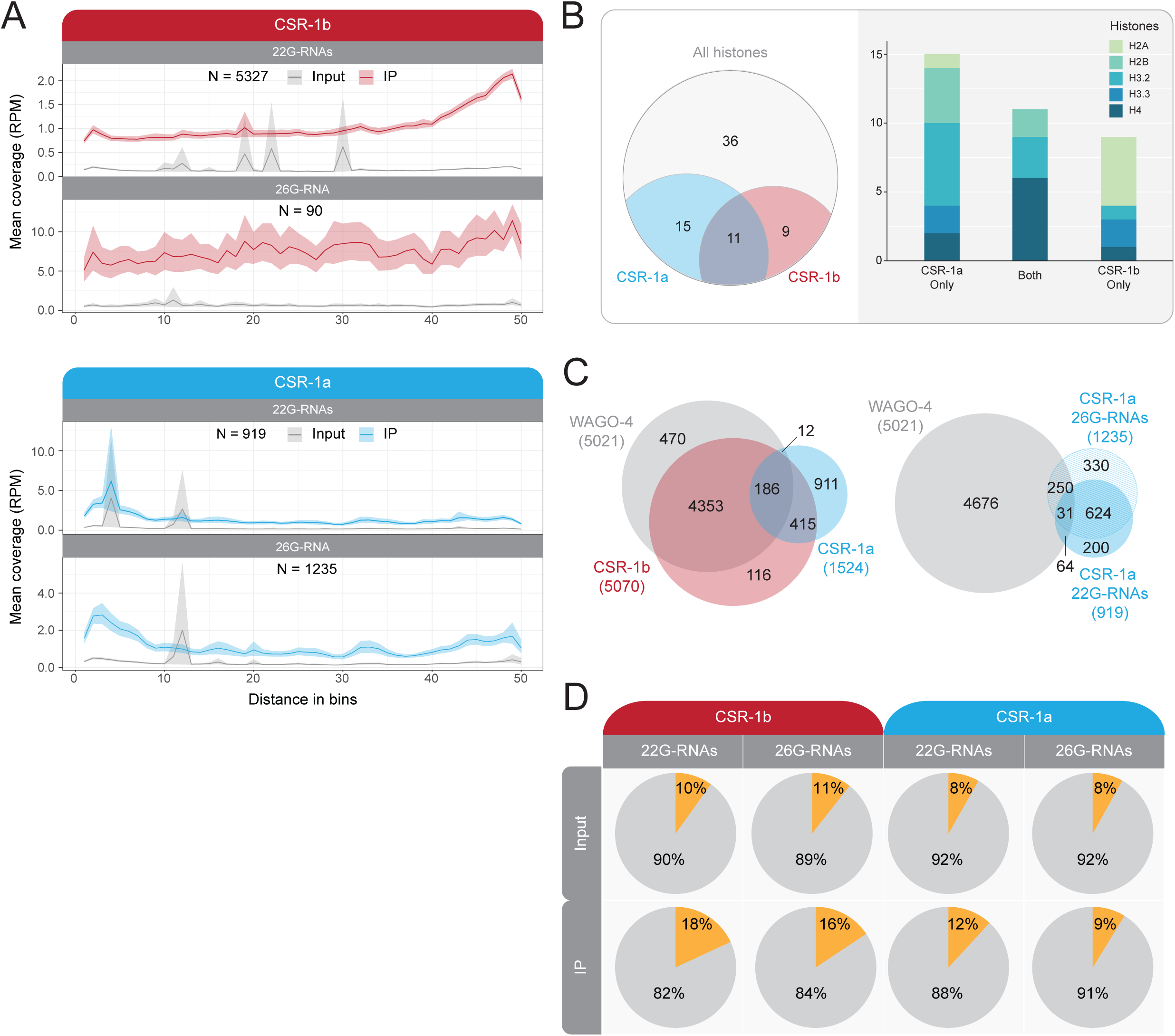
Additional comparisons of CSR-1b and CSR-1a sRNAs [Related to Fig. 3]. **A)** Metagene plot of the distribution of size-selected enriched sRNAs antisense to protein coding genes along the gene body for each CSR-1 isoform. N = number of genes, 22G = sRNAs in the range of 21-23nt, with no first nucleotide bias, 26G = sRNAs in the range of 25-27nt, with no first nucleotide bias. **B)** Venn diagram of *histone* genes enriched in the IPs for GFP::3xFLAG::CSR-1b and GFP::3xFLAG::CSR-1a, relative to all *histone* genes in *C. elegans*. **C)** Venn Diagrams of the overlap in targets from IPs of GFP::3xFLAG::CSR-1a, GFP::3xFLAG::CSR-1b, and GFP::3xFLAG::WAGO-4, using all sRNA sizes and no first nucleotide bias (left). GFP::3xFLAG::WAGO-4 all sRNAs targets overlapped with GFP::3xFLAG::CSR-1a 26G-RNA and CSR-1a 22G-RNA targets (right). **D)** Pie charts of percentage of total reads with U-tailing (gold) for Input and IP samples of GFP::3xFLAG::CSR-1b and GFP::3xFLAG::CSR-1a. 22G = sRNAs in the range of 21-23nt with no first nucleotide bias, 26G = sRNAs in the range of 25-27nt with no first nucleotide bias. Bar graph showing the distribution of types of histone genes enriched in each isoform IP.

**Supplemental Figure 8.**
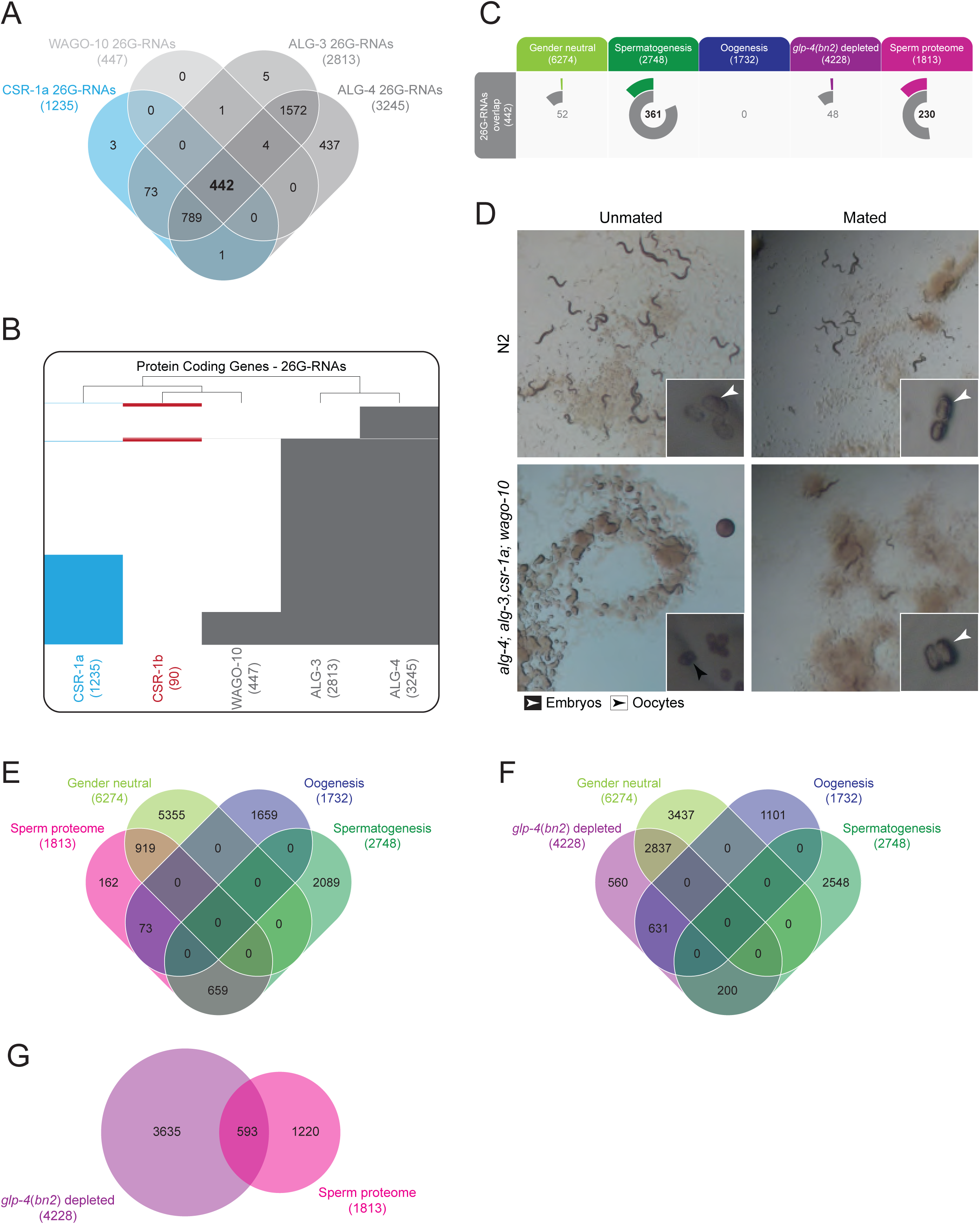
Additional analysis of spermatogenesis Argonautes [Related to Fig. 4]. **A)** Venn diagram depicting overlap in the protein coding gene targets of 26G-RNAs enriched in CSR-1a, ALG-3, ALG-4, and WAGO-10 IPs. A core set of 442 protein genes is shown in bold. **B)** Clustering diagram of genes targeted by 26G-RNAs (26G-sRNAs = sRNAs in the range of 25-27nt with no first nucleotide bias) enriched in IPs of each AGO. **C)** Venn-pie diagrams comparing groups of genes targeted by the core set of 442 genes targeted by all 26G-RNA binding AGOs (inner circle) with gene sets that define spermatogenesis-enriched transcripts, oogenesis-enriched transcripts, and gender neutral transcripts (enriched in both spermatogenesis and oogenesis) (Ortiz et al., 2014), germline expressed sRNAs (as determined by genes that are 2 fold or more depleted of sRNAs in *glp-4(bn2)* mutants) (Gu et al., 2009), and sperm proteome-enriched genes (Ma et al., 2014) (outer circle). The size of each color-coded circle represents the proportion of each sample that is shared between the two sets. Numbers in the center of the circle indicate the number of genes that overlap between the two data sets. Numbers in bold indicate statistically significant overlap as calculated by Fisher’s Exact Test, *p*<0.001. **D)** Images of wild-type worms and the quadruple spermatogenesis *ago* mutant with and without mating to wild-type males. Insets show magnification of embryos (white arrows) and unfertilized oocytes (black arrows). **E)** Venn diagram depicting overlap in germline gene sets (Ortiz et al., 2014) with sperm proteome-enriched genes (Ma et al., 2014). **F)** Venn diagram depicting overlap in germline gene sets with germline expressed sRNAs (as determined by genes that are 2 fold or more depleted of sRNAs in *glp-4(bn2)* mutants (Gu et al., 2009)). **G)** Venn diagram depicting overlap in sperm proteome-enriched genes (Ma et al., 2014) with germline expressed sRNAs (as determined by genes that are 2 fold or more depleted of sRNAs in *glp-4(bn2)* mutants (Gu et al., 2009)).

**Supplemental Table S1. Library statistics for all small RNA libraries used in this study.**

**Supplemental Table S2. CSR-1 isoform sRNA targets**

**Supplemental Table S3. CSR-1 isoform protein coding gene targets intersected with germline gene expression data**

**Supplemental Table S4. Spermatogenesis sRNA pathway protein coding gene target intersections**

**Supplemental Table S5. Gene Ontology (GO) analysis of spermatogenesis sRNA pathway protein coding gene targets**

**Supplemental Table S6. mRNA-seq differential expression analysis of *csr-1a(tor159)***

**Supplemental Table S7. Oligonucleotides and plasmids utilized in this study**

