## Supplementary material for "Two isoforms of the essential *C. elegans* Argonaute CSR-1 differentially regulate sperm and oocyte fertility through distinct small RNA classes": Key Resources Table

| **REAGENT or RESOURCE** | **SOURCE** | **IDENTIFIER** |
| --- | --- | --- |
| **Antibodies** | | |
| Rabbit anti-CSR-1 (total)  (peptide antigen: VDYNAPKDPEFRQKYPNLKFP) | Claycomb et al., 2009 | 49-C1 |
| Rabbit anti-CSR-1 (total)  (peptide antigen: QRCKDKGMHIGSYSMDQHNGERGSENFL) | Claycomb et al., 2009 | 49-C2 |
| Rabbit anti-CSR-1a  (peptide antigen: YQGKVKYRGGDRHDFNYEKTPTGSY) | This study | 55 |
| Rabbit anti-GFP | Torrey Pines | RRID:AB_2313770 |
| Mouse anti-alpha Tubulin | Sigma | RRID: AB_477593 |
| Monoclonal ANTI-FLAG M2 antibody | Sigma-Aldrich | RRID: AB_262044 |
| **Bacterial and Virus Strains** | | |
| *E. coli OP50* | Grown in house | N/A |
| **Chemicals, Peptides, and Recombinant Proteins** | | |
| cOmplete™, EDTA-free Protease Inhibitor Cocktail | Roche | Cat#4693132001 |
| Phosphatase Inhibitor Cocktail 2 | Sigma-Aldrich | Cat#P5726 |
| Phosphatase Inhibitor Cocktail 3 | Sigma-Aldrich | Cat#P0044 |
| SUPERase·In RNase Inhibitor | Invitrogen | Cat#AM2694 |
| GFP-Trap_MA Beads | ChromoTek | Cat#gtma-10 |
| Bolt LDS Sample Buffer (4X) | Invitrogen | Cat#B0007 |
| Bolt Sample Reducing Agent (10X) | Invitrogen | Cat#B0009 |
| Luminata Classico Western HRP Substrate | Millipore | Cat#WBLUC100 |
| RNA 5´ Polyphosphatase | Lucigen | Cat#RP8092H |
| Glycogen (5 mg/ml) | Ambion | Cat#AM9510 |
| TRI Reagent | Sigma-Aldrich | Cat#93289 |
| Ambion DNase I (RNase-free) | Ambion | Cat#AM2222 |
| UltraPure DNase/RNase-Free Distilled Water | Invitrogen | Cat#10977023 |
| **Critical Commercial Assays** | | |
| DC Protein Assay Kit | Bio-Rad | Cat#5000111 |
| NEBNext Ultra II Directional DNA library Prep Kit for Illumina | New England Biolabs | Cat#E7760S |
| NEBNext Poly(A) mRNA Magnetic Isolation Module | New England Biolabs | Cat#E7490S |
| NEBNext Multiplex Small RNA Library Prep Set for Illumina | New England Biolabs | Cat#E7300S |
| Gentra Puregene Tissue Kit | Qiagen | Cat#158689 |
| NEBNext DNA Library Prep Master Mix Set for Illumina | New England Biolabs | Cat3E6040 |
| NEBuilder HiFi DNA Assembly Kit | New England Biolabs | Cat#E2621 |
| **Deposited Data** | | |
| Small RNA Sequencing Libraries | This Study | GEO: GSE154678 |
| mRNA Sequencing Libraries | This Study | GEO: GSE154678 |
| *glp-4(bn2)* small RNA data | Gu et al., 2009 | GEO: GSE18215 |
| Genes expressed in the germline | Ortiz et al., 2014 | GEO: GSE57109 |
| Sperm Proteome Data | Ma et al., 2017 | http://159.226.118.206/miaolab/C.elegans%20data.htm. |
| Early Embryo and L4 expression data | Gerstein et al., 2010; Spencer et al., 2011 | SRX004866 and SRX008144. |
| WAGO-4 IP/Small RNA Seq | Xu et al., 2018 | GSE112475 |
| All custom code for this study | This study | www.github.com/ClaycombLab/Charlesworth_2020 |
| **Experimental Models: Cell Lines** | | |
| **Experimental Models: Organisms/Strains** |  |  |
| Wild-type *C. elegans*, Bristol Strain | CGC | N2 |
| *csr-1(tor67[gfp::3xflag::csr-1]) IV*  [made with CRISPR with SEC] | Ouyang et al., 2019 | JMC101 |
| *csr-1(tor160[csr-1 exon1::GFP::FLAG IV:7957568]) IV* [made with CRISPR with SEC] | This study | JMC151 |
| *csr-1(tor67[csr-1 exon2::GFP::FLAG IV:7958598]csr-1(nic361[G120*]) IV) IV*  [made with Co-CRISPR into JMC101] | This study | JMC164 |
| *csr-1(tm892) IV/nT1 [unc-?(n754) let-?] (IV;V)* | Yigit et al., 2006; Claycomb et al., 2009 | WM182 |
| *unc-119(ed3) III; mgTi13[rpl-28::gfp::csr-1b]; csr-1(tm892)* | This study | JMC160 |
| *unc-119(ed3) III; mgTi10[rpl-28::csr-1a exon1::gfp];csr-1(tm892)* | This study | JMC161 |
| *mgIs75[rpn-2::gfp] IV; mgTi14[pbs-5::mcherry]* | This study | GR3035 |
| *unc-119(ed3) III; mgTi10[rpl-28::csr-1a exon1::gfp]*  [Made with MiniMos] | This study | GR3036 |
| *unc-119(ed3) III; mgTi11[rpl-28::csr-1a exon1::gfp::csr-1]*  [made with MiniMos] | This study | GR3037 |
| *unc-119(ed3) III; mgTi12[rpl-28::csr-1a exon1::gfp::csr-1]*  *[Made with MiniMos* | This study | GR3038 |
| *csr-1(mg657[G91R]) IV* | This study | GR3039 |
| *unc-119(ed3) III; mgTi13[rpl-28::gfp::csr-1b]* | This study | GR3041 |
| *csr-1(mg660[G120*]) IV* | This study | GR3042 |
| *csr-1(mg661[G120*]) mgIs75[rpn-2::gfp] IV; mgTi14[pbs-5::mcherry]* | This study | GR3043 |
| *alg-4(tm1184) III; csr-1(tor67[csr-1 exon2::GFP::FLAG IV:7958598]csr-1(nic361[G120*]) IV) IV, alg-3(tm1155) IV; wago-10(tor133) V* | This study | JMC245 |
| *alg-3(tor141[alg-3::GFP::FLAG]) IV* | This study | JMC205 |
| *alg-4(tor143[alg-4::GFP::FLAG]) III* | This study | JMC207 |
| *wago-10(tor127[wago-10::GFP::FLAG]) V* | This study | JMC233 |
| *wago-4(tor117[wago-4::GFP::FLAG]) II* | Lev et al., 2019 | JMC223 |
| *pha-1(e2123)* | Schnabel and Schnabel, 1990 | GE24 |
| *znfx-1(gg634[HA::tagRFP::znfx-1]) II.* | Wan et al., 2018 | YY1446 |
| **Oligonucleotides** | **Related plasmid no.** | **Identifier** |
| See Supplemental Table S7 |  |  |
| **Recombinant DNA** | | |
| See Supplemental Table S7 |  |  |
| **Software and Algorithms** | | |
| FastQC | Andrews, 2010 | RRID:SCR_014583 |
| Cutadapt | Martin, 2011 | RRID:SCR_011841 |
| STAR | Dobin et al., 2013 | RRID:SCR_011841 |
| sRNA Counting Scripts | This study | www.github.com/ClaycombLab/Charlesworth_2020 |
| Enrichment Script | This study | www.github.com/ClaycombLab/Charlesworth_2020 |
| ggplot2 | Wickham, 2016 | https://ggplot2.tidyverse.org |
| VennDiagram Package | Chen and Boutros, 2018 | https://CRAN.R-project.org/package=VennDiagram |
| VennDetail Package | Guo and McGregor 2020 | https://github.com/guokai8/VennDetail. |
| Metagene2 | Fournier et al., 2020 | https://github.com/ArnaudDroitLab/metagene2. |
| pheatmap | Kolde et al., 2019 | https://CRAN.R-project.org/package=pheatmap |
| DeSeq2 | Love et al., 2014 | RRID:SCR_015687 |
| EnhancedVolcano  v. 1.4.0 | Blighe et al., 2020 | https://github.com/kevinblighe/EnhancedVolcano |
| GenomicFeatures | Lawrence et al., 2013 | doi:10.1371/journal.pcbi.1003118 |
| R | N/A | https://www.R-project.org |
| R Studio | RStudio Team (2020) | <http://www.rstudio.com/> |
| InterMine/ WormMine | Smith et al., 2012; Kalderimis et al., 2014 | http://intermine.wormbase.org/tools/wormmine/ |
| WormExp | Yang et al., 2016 | http://wormexp.zoologie.uni-kiel.de/wormexp/ |
| Wormbase | N/A | http://www.wormbase.org |
| g:Profiler | N/A | https://biit.cs.ut.ee/gprofiler/gost |
| GraphPad Prism | N/A | https://www.graphpad.com/ |
| NIS Elements AR Software | N/A | https://www.microscope.healthcare.nikon.com/ |
| Fiji | N/A | https://fiji.sc/ |
| **Other** |  |  |
| Bolt 4 to 12%, Bis Tris 1.0 mm Mini Protein Gel | Invitrogen | Cat#NW04120BOX |
